# Dynamic control of cortical head-direction signal by angular velocity

**DOI:** 10.1101/730374

**Authors:** Arseny Finkelstein, Hervé Rouault, Sandro Romani, Nachum Ulanovsky

## Abstract

The sense of direction requires accurate tracking of head direction at different turning-velocities, yet it remains unclear how this is achieved in the mammalian brain. Here we recorded head-direction cells in bat dorsal presubiculum and found that, surprisingly, the head-direction signal in this cortical region was dynamically controlled by angular velocity. In most neurons, a sharp head-direction tuning emerged at some angular velocity, but was absent at other velocities – resulting in a 4-fold increase in head-direction cell abundance. The head-direction tuning changed as a function of angular velocity primarily via a redistribution of spikes between the neuron’s preferred and null directions – while keeping the average firing-rate constant. These results could not be explained by existing ‘ring-attractor’ models of the head-direction system. We propose a novel recurrent network model that accounts for the observed dynamics of head-direction cells. This model predicts that the new classes of cells we found can improve the sensitivity of the head-direction system to directional sensory cues, and support angular-velocity integration.

## Introduction

Spatial orientation is one of the most fundamental forms of behavior^1^. Head-direction (HD) cells, originally discovered in the rat dorsal presubiculum, are neurons that are tuned to head azimuth – with each cell tuned to a specific direction, together spanning the entire 360° azimuth range^2, 3^. Neurons with HD tuning appear during development before all the other spatially-tuned cells^4, 5^, and are believed to provide the animal with a sense of direction by acting as a neural analogue of a compass^6–10^.

The generation of HD tuning was suggested to depend on a combination of: (i) a self-motion signal that relies mostly on the vestibular system, and is processed in subcortical areas^11–14^; (ii) information about head-orientation with respect to external landmarks, which comes from visual areas^15–18^; and (iii) internal network dynamics that are independent of external sensory inputs^19–25^. Network models of HD cells typically include a ring-attractor network, in which neurons of similar preferred-orientation share excitatory connections, while neurons with dissimilar tuning inhibit each other^22–24, 26^. This structure of local-excitation and long-range inhibition^27^ can lead to the formation of a stable ‘bump’ of activity, representing the current direction of the neural compass.

A critical feature of the HD circuitry is the ability to update the HD cell activity during head turns – a feature that was suggested to result from angular-velocity integration^7, 28–30^. In network models, this was implemented by a 3-ring architecture with asymmetric connectivity (Fig. 1a, see legend for details of the connectivity), which allows integrating a putative angular-velocity input and shifting the bump of activity along the ring attractor by an angle corresponding to the actual head turn^10, 22–24, 26, 31, 32^. Importantly, for angular-velocity integration, these models critically rely on the existence of neurons that conjunctively encode head direction and angular velocity (Fig. 1b,c). Recently, experiments in the fly central complex have shown that indeed a ring attractor-like network organization containing such conjunctive neurons is responsible for maintaining and updating the HD signal^33–35^. To date, however, in the mammalian brain only a tiny fraction of HD cells was reported to have conjunctive tuning to angular-velocity^12, 14^ These neurons were found exclusively in subcortical regions along the vestibular pathway, despite the fact that HD cells are distributed across both subcortical and cortical regions^30^.

**Fig. 1.**
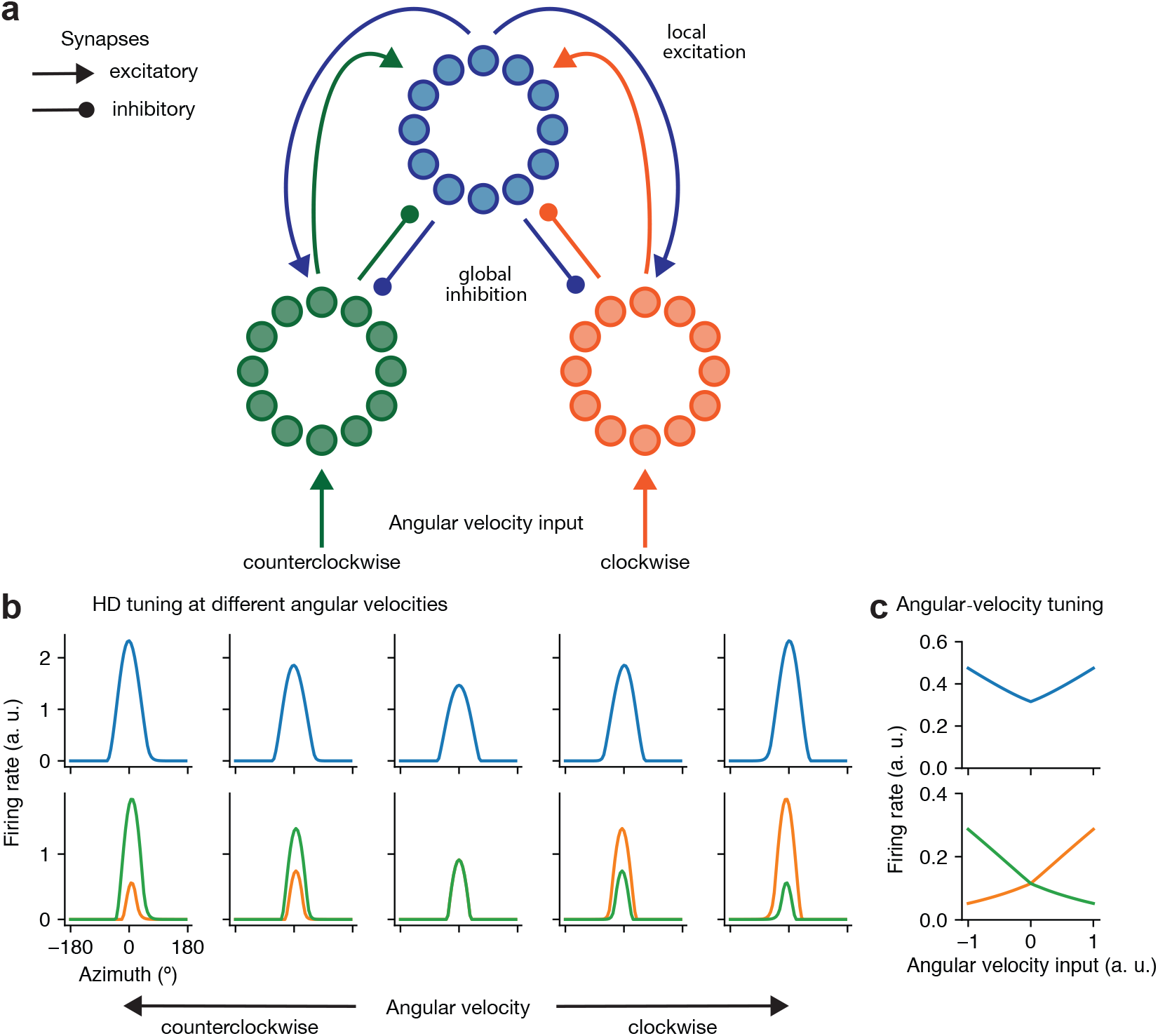
Classical 3-ring attractor model of head-direction cells. **a**, HD cell activity has been classically modeled with ‘ring attractor’ networks^10, 22–24, 26, 29–32, 35^. Classical ring-attractor models contain neurons that are recurrently connected with connectivity patterns that match the topology of a ring. In order to update the position of the activity-bump following head-movement, using angular-velocity integration, the network includes two additional populations of neurons (‘left’ and ‘right’ rings, respectively green and orange). These populations are reciprocally connected with the first one (‘center’ ring, blue) and form activity bumps, but they synapse onto the center ring asymmetrically: the connections from the left ring are shifted counterclockwise, while those from the right ring are shifted clockwise. The left and right rings also receive angular-velocity inputs that are responsible for turning the position of the bump in the central ring. For instance, during clockwise (CW) turns, there is a stronger input to the right ring, which would pull the central bump in the clockwise direction. Conversely, during counterclockwise (CCW) turns, there is a stronger input to the left ring, and the bump would move in the counterclockwise direction. **b**, Simulated HD-tuning curves at different angular velocities for neurons on different rings (blue – central ring, green – left ring, orange – right ring). The central column corresponds to zero angular-velocity (absence of head-turns). The two left columns correspond to increasing counterclockwise velocity, and the two right columns correspond to increasing clockwise velocity. Note the symmetric modulation (scaling) of HD tuning as a function of angular velocity for neurons on the central ring (blue: top row), and the asymmetric modulation for neurons on the left and right rings (green and orange: bottom row). **c**, Angular-velocity tuning. Classical 3-ring models, like the one we considered here, predict that modulation of HD tuning by angular velocity should be accompanied by an overall change in the firing-rate of the cell. Our experimental data was at odds with this prediction (**Fig. 3,4**) – which requires a revision of the classical models.

We, therefore, hypothesized that cortical regions might also harbor a representation of a conjunctive HD × angular-velocity signal – perhaps with different properties than in subcortical regions. In particular, in addition to vestibular inputs, cortical circuits may extract information about head-movement from multiple other sources, including visual, proprioceptive, and motor efference-copy signals^9^. We focused here on the dorsal presubiculum, which is the first cortical region to receive both HD inputs from the vestibular pathway and visual inputs, and is also the main gateway of HD input into the medial entorhinal cortex^16, 36–39^. Electrophysiological recording from rat dorsal presubiculum revealed that the mean firing-rate of HD cells was unaffected by angular velocity^40^. This has led to previous suggestions that the generation and update of the HD signal via angular-velocity integration happens subcortically^30^, whereas cortical circuits inherit a pure representation of HD, which is not affected by angular velocity^16^. By contrast, we previously observed modulation of three-dimensional (3D) HD tuning (to azimuth, pitch, and roll of the head) by angular velocity, in the dorsal presubiculum of bats^41, 42^. This prompted us to examine in more detail the effects of angular velocity on the azimuthal HD tuning in this cortical region in bats.

Here we found that contrary to the common view, the tuning of most HD neurons in the dorsal presubiculum changes profoundly as a function of angular velocity. Most neurons increased their directional tuning at faster angular velocities, often showing preference to either clockwise or counterclockwise turning directions – as predicted by angular-velocity integration models (Fig. 1b,c), but which so far were attributed only to subcortical regions. Surprisingly, for most cells this modulation was not accompanied by changes in the mean firing-rate of the neuron over all HD angles (we will refer to change in the mean-firing rate of a neuron as gain). Instead, we observed a change in the modulation depth of the HD-tuning curve (scaling) as function of angular velocity, which occurred via a redistribution of spikes at the null versus preferred directions – demonstrating a new form of a conjunctive tuning (see Supplementary Fig. 1 for a schematic illustration of HD-tuning scaling via gain or redistribution of spikes). Further, some neurons that were completely un-tuned to HD when measured in the conventional manner (averaging over all angular velocities), were found to exhibit strong directional tuning at specific angular velocities. These dynamic effects of angular velocity on the HD tuning are difficult to explain by existing ring attractor models of HD cells (Fig. 1). However, we show that these novel non-classical phenomena can be explained by extending the architecture and components of classical network models of HD tuning – supporting a different theoretical approach for generating and updating the HD signal at the cortical level.

## Results

### Emergence of head-direction tuning at different angular velocities

We recorded the activity of individual neurons in the dorsal presubiculum of Egyptian fruit bats that crawled on the horizontal surface of an arena in search for food^41^. We analyzed here the azimuthal HD and angular-velocity (AV) tuning of 346 cells×sessions that met inclusion criteria for the number of spikes and behavioral sampling of angular velocity (Methods).

For each neuron, we first computed the conventional HD tuning-curve based on the behavioral and neural data collected over the entire session (Fig. 2a, column i). Across all recorded cells, 17.1% were significantly directionally-tuned according to their conventional HD-tuning curves (Methods), and we will refer to them as ‘conventional HD cells’.

**Fig. 2.**
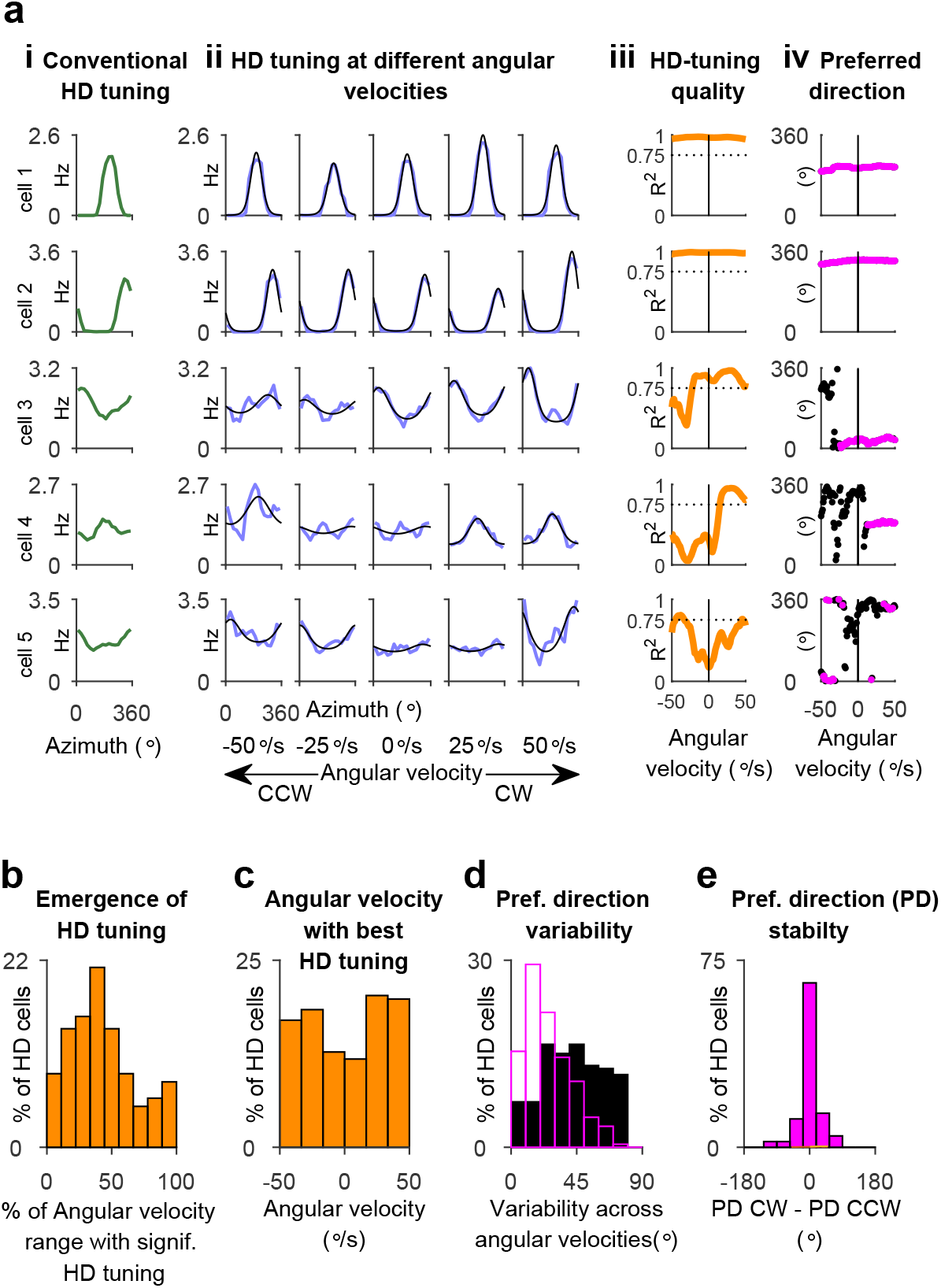
Head-direction tuning is controlled by angular velocity. **a**, Example cells (rows) showing the effects of angular velocity on the head-direction (HD) tuning curve. Columns: *i*, Conventional HD tuning computed for the entire session. *ii*, HD tuning curves (blue) computed for different nonoverlapping angular-velocity bins, overlaid with von Mises fits (black). Numbers below each column indicate the center of the angular-velocity bin; CW, clockwise; CCW, counterclockwise rotations of the head. *iii*, Quality of HD tuning (assessed by goodness-of-fit, *R*^2^, of the Von Mises function to the experimental tuning), as a function of angular velocity. *iv*, Preferred direction as a function of angular velocity, for significantly-tuned angular-velocity bins (magenta) and untuned angular-velocity bins (black). In *iii* and *iv*, we used a sliding window of 25°/s that was moved in steps of 1°/s. **b-e**, Population analyses for the 212 cells×sessions which exhibited significant directional tuning in at least some angular velocity. **b**, Percentage of angular-velocity range for which neurons exhibited significant directional tuning. **c**, Distribution of angular velocities for which the neurons had the best HD tuning (best fit to von Mises function). **d**, Distribution of the variability in the preferred direction across tuned angular-velocity bins (magenta) and untuned angular-velocity bins (black), for all the cells with HD tuning. **e**, Distribution of the differences in preferred direction (PD) between HD tuning curves for CW versus CCW angular-velocity bins. The comparison was done for tuned angular-velocity bins only (Methods).

We next examined the effects of angular velocity on the HD tuning, by splitting the data and computing the HD tuning at different angular velocities (Fig. 2a, column ii; see distribution of angular velocities in Supplementary Fig. 2a, column v). We found conventional HD cells that were tuned across the entire sampled angular-velocity range (Fig. 2a, columns i-ii, cells 1–2). Surprisingly, in many neurons that were *not* categorized as conventional HD cells, a significant directional tuning emerged at a certain angular-velocity range (Fig. 2a, compare tuning in columns i and ii for cells 4–5; and see Supplementary Fig. 3 for additional examples of neurons which did *not* exhibit classical HD tuning but where HD tuning emerged at specific angular velocities). We will refer to neurons that were tuned to HD, either at some or at all angular velocities, as ‘HD cells’. This suggests that the conventional HD-tuning curve does not provide an adequate depiction of directionality – because it combines both tuned and untuned epochs. In fact, 61.3% of all recorded cells exhibited significant HD tuning in at least some of the angular-velocity range tested (Methods) – almost 4 times more than we and others found by conventional analysis of HD tuning in the dorsal presubiculum of rats and bats^2, 41, 43^.

To better quantify the effects of angular velocity, we analyzed the change in HD-tuning quality as a function of angular velocity, by assessing the goodness-of-fit of the observed tuning to a circular unimodal distribution (Fig. 2a, column iii; Methods). We found that in the majority of recorded neurons, a unimodal directional tuning emerged at some angular velocities, with most cells being tuned in < 50% of the angular-velocity range (Fig. 2b). There was no relationship between the quality of the tuning in a particular angular-velocity bin and the time spent at this angular velocity (Supplementary Fig. 2a – compare column ii and v; and Supplementary Fig. 2c). This indicates that loss of HD tuning at some angular velocities does not result from an insufficient sampling of the directional tuning curve. Angular velocities at which different cells were best tuned to head direction were found to span the entire velocity-range sampled (Fig. 2c), with transitions from ‘tuned’ to ‘untuned’ states occurring at different angular-velocity ranges for different cells (Supplementary Fig. 3; and see Methods). Overall, there was a slightly higher propensity of cells to be directionally-tuned at faster angular-velocities (Fig. 2c: note the notch around zero angular-velocity; χ^2^ test for non-uniformity: *P* = 0.05).

We next examined whether the HD tuning measured at different angular velocities had a stable directional preference. To test this, we computed the preferred direction as a function of angular velocity and found that for angular velocities for which the neuron exhibited HD tuning, the preferred direction was stable (Fig. 2a, column iv: note that the magenta dots, which denote significantly-tuned bins [Methods], had a stable preferred direction). By contrast, degradation of HD tuning was associated with irregular fluctuations of the preferred direction (Fig. 2b, column iv: note the black dots, which denote untuned bins [Methods]). At the population level, the HD tuning was rather stable as a function of angular velocity – with a much smaller variability of preferred-directions across tuned angular-velocity bins (Fig. 2d, magenta), as compared to untuned bins (Fig. 2d, black). The stability of HD tuning as a function of angular velocity was also confirmed by computing the tuning separately for clockwise and counterclockwise head-turns, and comparing the preferred firing-direction for both turning directions (Fig. 2e, note the narrow distribution around 0° – indicating no change in the preferred direction of the neurons between head-turns in opposite directions; only HD cells that were tuned for both turning-directions were used in this analysis: see Methods). This suggests that HD cells maintain the same directional preference during head turns at some angular velocities – but for other angular velocities, the directional tuning can disappear entirely (see Fig. 2a, cells 3–5).

### Scaling of head-direction tuning by angular velocity

We next examined whether angular velocity not only controls categorically if HD tuning would emerge or disappear, but may also gradually scale the modulation depth of the directional tuning (where modulation-depth quantifies the firing-rate at the preferred direction relative to the null direction; Methods; see illustration in Supplementary Fig. 1a). We defined *scaling* of the HD tuning as a change in modulation depth as a function of angular velocity (Fig. 3a-b, column iii – HD-tuning scaling; Methods). Almost all HD cells had significant HD-tuning scaling by angular velocity (Fig. 3a-b, columns ii,iii), constituting 60.4% of all recorded cells (Fig. 3c – blue circle). Such scaling of HD-tuning could not be explained by random fluctuations in firing rate and inhomogeneous sampling (Supplementary Fig. 2a, d). Interestingly, for most cells, the HD tuning-scaling was largest at high angular velocities (Fig. 3d, note the U-shaped distribution; χ^2^ test for nonuniformity: *P* < 0.001). Across the population, HD-tuning scaling often showed many-fold changes in modulation depth as a function of angular velocity (Fig. 3e) – the first demonstration of such scaling in cortical circuits.

**Fig. 3.**
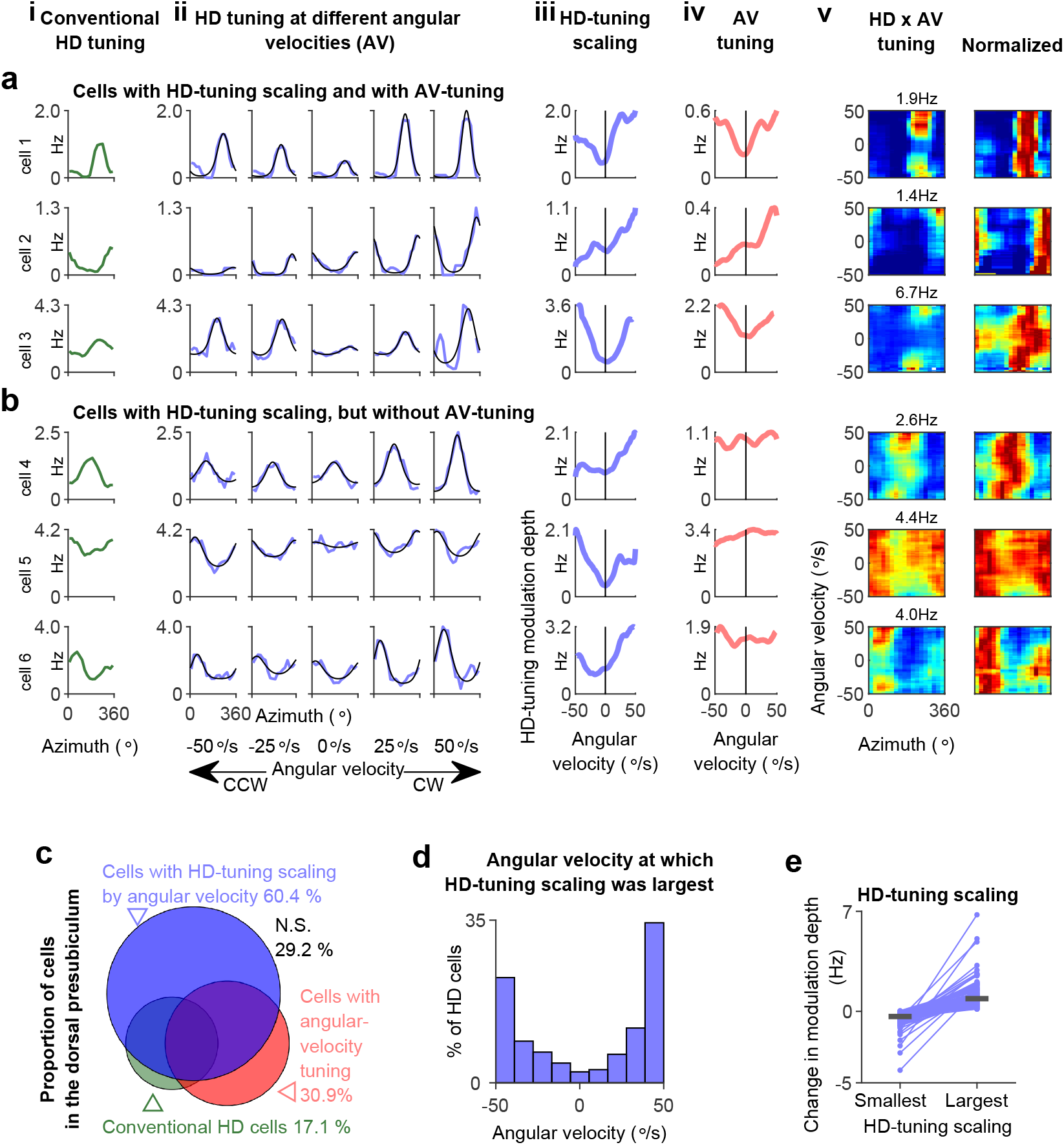
Scaling of head-direction tuning by angular velocity. **a-b**, Example cells (rows) with HD-tuning scaling by angular velocity. Columns: *i*, Conventional HD tuning computed for the entire session. *ii*, HD-tuning curves (blue) computed for epochs with different angular velocities, overlaid with von Mises fits (black). *iii*, HD-tuning scaling (i.e. changes in the modulation depth of HD tuning curve) as a function of angular velocity. *iv*, Angular-velocity tuning. *v*, Firing-rate map of HD tuning computed for different angular velocities (rows), color-scaled to the highest HD peak firing rate across all angular velocities (left), or with each HD-tuning curve (each row) normalized to its own peak (right). **c**, Venn diagram showing the proportion of cells×sessions in the dorsal presubiculum exhibiting significant HD-tuning scaling or AV tuning. Note that most cells with significant HD tuning-scaling (blue circle) were *not* classified as HD cells according to the conventional analysis (green), and did *not* exhibit significant AV tuning (red). **d**, Distribution of angular velocities in which the modulation depth of the HD tuning was maximal, plotted for all cells with significant HD-tuning (*n* = 212 cells×sessions). **e**, Change in HD-tuning modulation-depth (for smallest or largest HD-tuning scaling), relative to the modulation depth of the conventional HD tuning, for same cells as in d.

### Conjunctive head-direction×angular-velocity tuning via gain or redistribution of spikes

HD tuning by angular velocity could simply reflect a tuning to angular velocity – i.e., a change in firing-rate as a function of angular velocity, regardless of the HD tuning (Methods). Indeed, we found that 30.9% of all recorded neurons had a significant AV tuning (Fig. 3c, red circle; Methods). Most of these cells were also tuned conjunctively to HD and showed HD-tuning scaling by angular velocity (Fig. 3a, columns iii-v; 21.4% of all recorded neurons; Fig. 3c, the intersection of red and blue circle). Surprisingly, however, a large fraction of neurons with HD-tuning scaling by angular velocity were *not* tuned to angular-velocity *per se* (Fig. 3b, columns iii-v; 39.0% of all recorded neurons; Fig. 3c, the part of the blue circle that does not intersect with the red circle).

To further characterize the relationship between HD-tuning scaling and AV tuning, we grouped all cells with HD-tuning into classes (Methods), based on the shape of their HD-tuning scaling (Fig. 4a, *x*-axis: blue icons) and on the shape of their AV-tuning (Fig. 4a, *y*-axis: peach-colored icons). We found neurons that scaled-up their HD tuning symmetrically for faster turns in both turning directions (column 1), or asymmetrically for clockwise (column 2) or counterclockwise directions (column 3). Analogously we found neurons with symmetric or asymmetric AV tuning (Fig. 4a, rows), or no AV tuning at all (second top row).

**Fig. 4.**
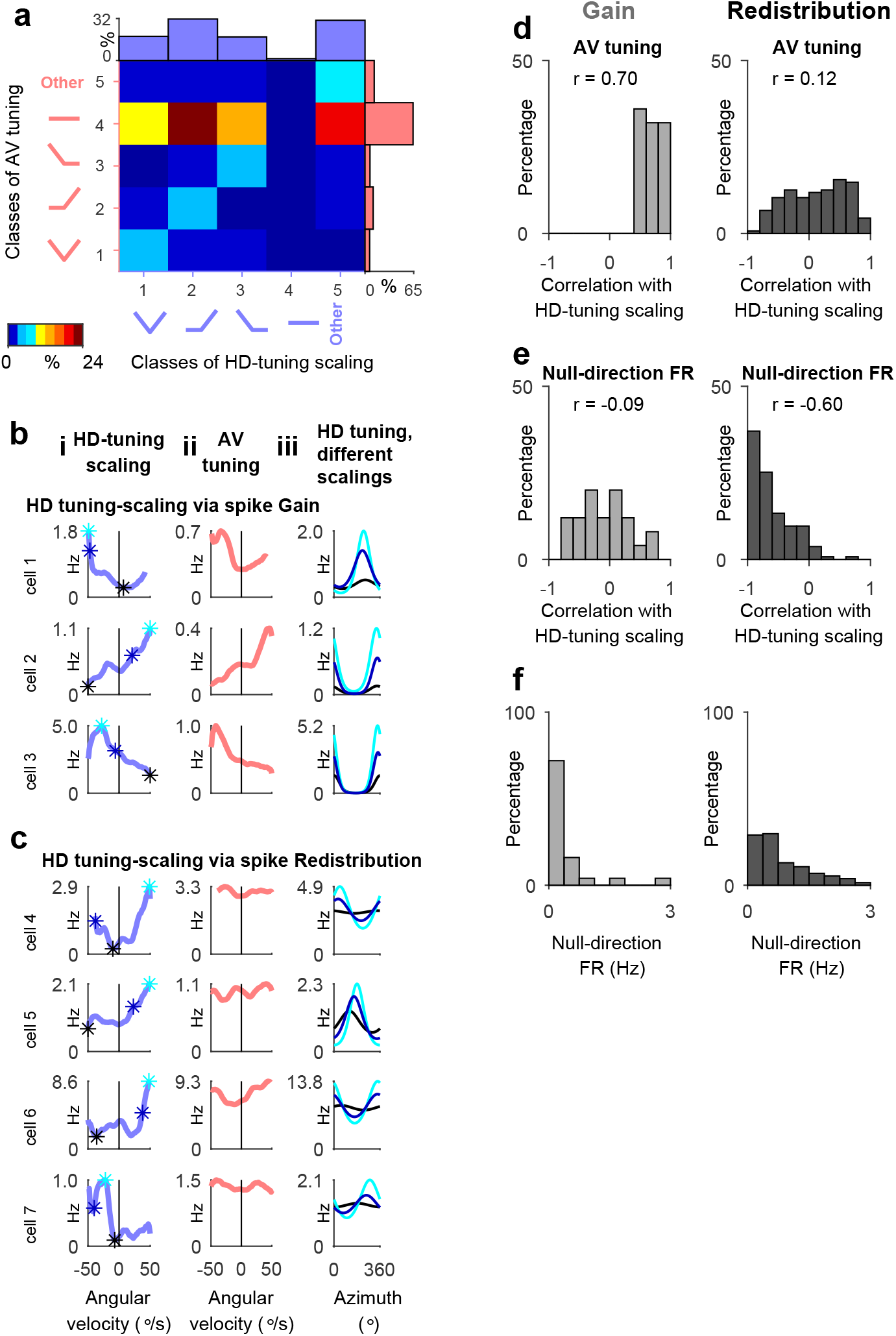
Conjunctive tuning to head direction and angular velocity involves two mechanisms: spike gain or spike redistribution. **a**, Classification of HD cells according to shapes of HD-tuning scaling and angular-velocity tuning. Percentage of HD cells grouped by different classes, based on the shape of HD-tuning scaling (columns) and the shape of the AV-tuning (rows); shown also are the marginal distributions of proportions of cells for the different classes. Cells with HD-tuning scaling but no AV tuning (4^th^ row of the matrix) were classified as ‘*redistribution*’ cells. Cells with HD-tuning scaling shape that mirrored the shape of the AV tuning (the bins on the diagonal of the matrix) were classified as ‘*gain*’ cells. **b-c**, Example cells (rows) with HD-tuning scaling via spike-gain (**b**) or spike-redistribution (**c**). Columns: *i*, HD-tuning scaling as a function of angular velocity. *ii*, Angular-velocity tuning. Note that the shape of AV tuning matched the shape of HD-tuning scaling for gain cells (b), but not for redistribution cells (c). *iii*, Left, Von-Mises fits of HD tuning computed for angular-velocity bins with largest (cyan), intermediate (blue), or smallest (black) HD-tuning scaling values; the corresponding angular-velocity bins are marked by asterisks in column *i*, with matching colors. **d**, Distribution of Pearson correlation coefficients between the shape of HD-tuning scaling and shape of AV-tuning plotted for gain cells (left) and redistribution cells (right). **e**, Same as in d, with correlation computed between the shape of HD-tuning scaling and the shape of null-direction firing-rate modulation as a function of angular-velocity. Note that in d-e, the distributions have opposite tendencies for the two groups of cells, indicating that changes in HD-tuning scaling for gain cells versus redistribution cells resulted from two distinct mechanisms. **f**, Distributions of firing-rates at the null direction for gain cells (left) and redistribution cells (right).

For a subset of neurons with conjunctive HD x AV tuning, the shape of the HD-tuning scaling by angular-velocity reflected the shape of the AV tuning (e.g. Fig. 4b, note similarity between the shape of curves in columns i and ii; see also the prominent diagonal band in the matrix in Fig. 4a). For these neurons, HD-tuning scaling by angular velocity was associated with a selective gain of spikes at the preferred direction, and we therefore termed these neurons *gain cells*. Gain cells comprised 7.2% of all recorded neurons and 11.8% of the HD cells (Methods). The existence of such cells with conjunctive HD x AV tuning via gain of spikes is predicted by classical 3-rings models (Fig. 1b-c, see legend for detailed explanation), but was not observed in the mammalian cortex so far.

However, most neurons exhibited HD-tuning scaling without AV tuning (manifested by the prevalence of cells in the 4th row in the matrix in Fig. 4a). In these cells, HD-tuning scaling by angular velocity occurred *without* a net change in spike count– and we therefore termed these neurons *redistribution cells* (e.g. Fig. 4c, compare columns i-ii). Neurons with HD-tuning scaling shape that did not fit into our classification scheme also mostly showed spike redistribution (Fig. 4a, row 4 column 5 [‘other’]). Redistribution cells comprised 39.0 % of all recorded cells and 63.7% of HD cells (Methods). Thus, most HD cells in the bat dorsal presubiculum did *not* encode angular velocity by net changes in average firing-rates of individual cells. Instead, they were conjunctively tuned to angular velocity indirectly, by HD-tuning scaling via redistribution of spikes (see Supplementary Fig. 1b for a schematic illustration of the gain and redistribution effects).

### Properties of Gain and Redistribution cells

The shape of HD-tuning scaling of both redistribution cells and gain cells was stable (Supplementary Fig. 4), indicating that HD cells had a consistent angular-velocity modulation. Interestingly, simultaneously-recorded HD cells showed a large heterogeneity in the shape of their HD tuning-scaling (Supplementary Fig. 5a). Specifically, we did not find systematic significant correlations in the shape of HD-tuning scaling of pairs of simultaneously-recorded cells (see Supplementary Fig. 5d; average Pearson correlation between HD-tuning scaling of simultaneously recorded cells: *r* = 0.08). Taken together, this suggests that HD-tuning scaling is stable across time but not necessarily shared by all cells in the HD network.

To further analyze the difference between redistribution cells and gain cells, we computed the correlation between the shape of HD-tuning scaling and the shape of AV tuning (Fig. 4d). Consistent with our definitions of gain and redistribution cells (Methods), we found a strong positive correlation between the two for gain cells – but not for redistribution cells (compare the two histograms in Fig. 4d). We then computed for each neuron the correlation between the shape of the HD-tuning scaling as a function of AV and the shape of null-direction firing-rate modulation as a function of AV (Fig. 4e). Importantly, we found that for redistribution cells, HD-tuning scaling was significantly *anti*-correlated with null-direction firing-rate – but this was not the case for gain cells (compare histograms in Fig. 4e).

This demonstrates that for redistribution cells, but not for gain cells, the increase in directional tuning at specific angular velocities involved a selective suppression of the firing rate at the non-preferred direction (Fig. 4b-c, column iii: note that null-direction firing rate for tuning with maximal modulation depth [cyan line] was lower than for tuning with intermediate [blue line] and minimal modulation depth [black line] – but only for the redistribution cells and not for the gain cells). In other words, redistribution cells maintained a constant mean firing-rate as a function of angular velocity, by simultaneously increasing the firing-rate at the preferred direction and correspondingly decreasing the firing-rate at the null direction. This dichotomy (Fig. 4b,c,e) further supports the notion that conjunctive tuning to head direction and angular velocity could arise from two distinct mechanisms – involving a selective gain of spikes or a balanced redistribution of spikes.

### Six-ring attractor network model of head-direction tuning

We next asked whether the existing network models of HD cells could capture these dynamic modulations of HD tuning by angular velocity. To this end, we considered the classical 3-ring network – an architecture used in many models of HD cells^23, 24, 26, 32^ (Fig. 1a). Simulated neurons in this model showed HD-tuning scaling by angular velocity (Fig. 1b). Importantly, in this model, HD-tuning scaling was always accompanied by a corresponding AV-tuning (Fig. 1c) – as in gain cells that we found experimentally. However, the presence of redistributions cells, for which HD-tuning scaling is not accompanied by AV tuning, could not be explained by existing models of HD cells. We therefore reasoned that additional cell populations might be required to account for the experimentally observed HD-tuning scaling via redistribution of spikes.

In most existing models, the relationship between input current and firing-rate is such that neurons are inactive when their inputs are below a threshold, and their activity grows monotonically with the magnitude of supra-threshold inputs. This ‘non-linearity’ allows the formation of a persistent and localized bump of activity on the ring for certain parameters of the connectivity-strength among neurons^44–46^. We found experimentally that cells which scaled via gain of spikes, and could be explained by the classical models, had a negligible firing-rate at their null-direction (Fig. 4f, left) – suggesting that they are inactive (subthreshold) at the null direction and become active (suprathreshold) at the preferred direction. By contrast, cells that redistributed their firing-rate as a function of angular velocity exhibited some baseline firing rate at their null-direction (Fig. 4f, right) – indicating that they primarily operated above the spiking-threshold. This suggests that redistribution cells might receive a constant input that maintains them above threshold.

To account for the redistribution phenomenon, we started with the classical 3-ring network architecture (Fig. 1a) – which is solely composed of neurons operating both in sub- and supra-threshold regimes (we would refer to these neurons as ‘*classical*’). We then added to this classical 3-ring network 3 additional populations of neurons: these 3 new populations were also organized in three rings similarly to the classical model (Fig. 5a; Methods). Additionally, they received a strong uniform external excitatory input, and a strong recurrent inhibitory feedback – and therefore we refer to these neurons as ‘*balanced*’ (Fig. 5b: classical neurons – solid lines, balanced neurons – dashed lines). These inputs clamped the mean firing-rate of these additional neurons to a constant supra-threshold value. When the local recurrent excitation between balanced neurons was sufficiently weak, most of these neurons were functioning above threshold. We found that these balanced neurons (although operating in a supra-threshold regime) could be recurrently coupled to classical neurons (operating in both sub- and supra-threshold regimes), to form a 6-ring network (see Fig. 5a-b and Methods for details of the coupling). It is possible to achieve this coupling without destroying the existence of a localized bump of activity (Supplementary Fig. 6; see Methods for the coupling constants), while still allowing velocity integration by the network (Supplementary Fig. 7).

**Fig. 5.**
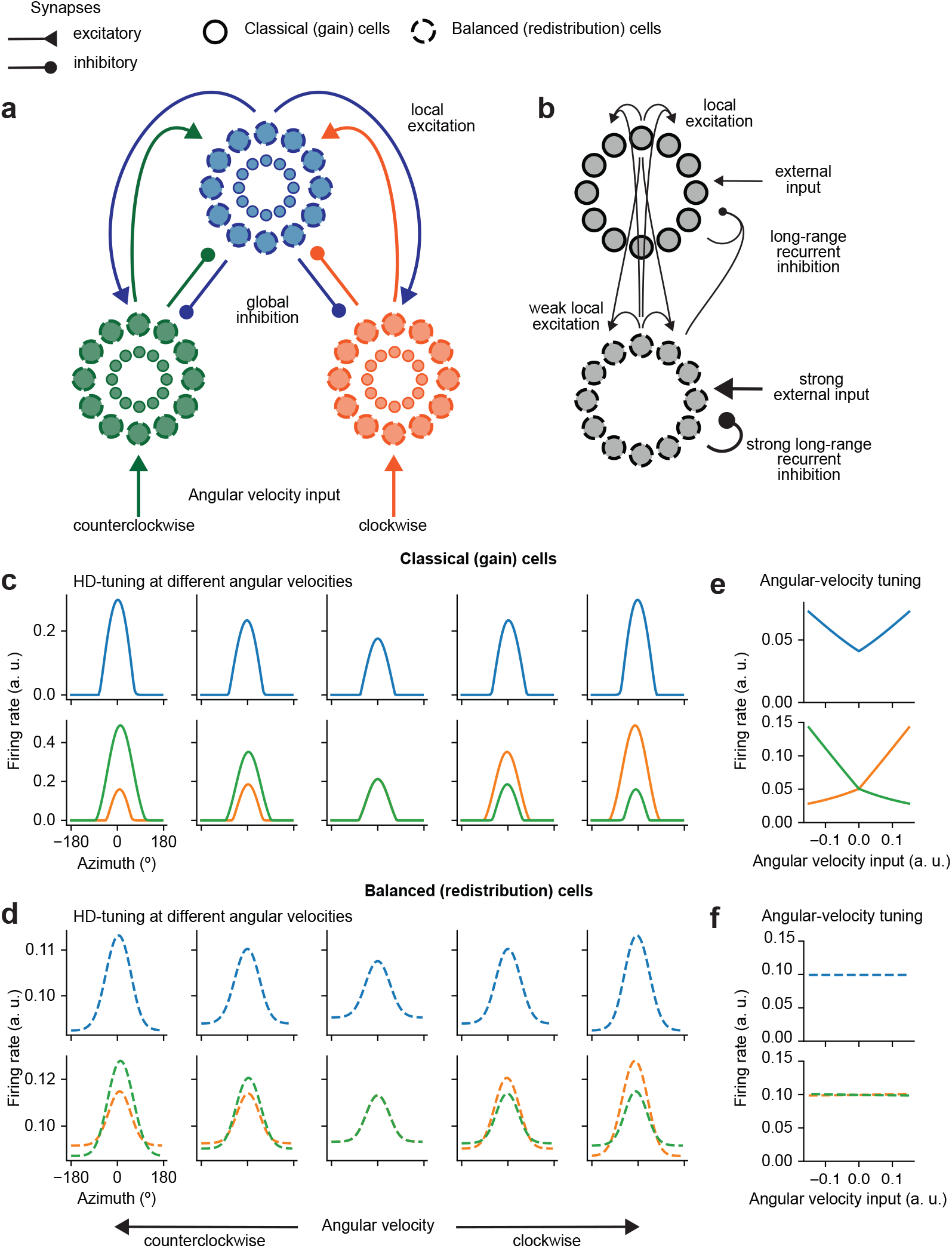
A 6-ring attractor model explains the experimentally observed modulations of the head-direction tuning by angular velocity. **a**, Schematics of the 6-ring model (see Methods for the numerical implementation) that consists of a total of six interconnected rings. Classical neurons (solid lines), balanced neurons (dashed). Blue rings correspond to the central rings that do not receive angular-velocity inputs. The side rings (orange and green) receive respectively the clockwise and the counterclockwise angular-velocity inputs, and excite the central rings with clockwise and counterclockwise shifts. **b**, Schematics of coupling between classical and balanced neurons in the 6-ring model, for each pair of rings (e.g. coupling between the central rings). Balanced neurons receive a strong feed-forward excitatory input compensated by a strong recurrent inhibitory feedback that controls their mean-firing activity. The other interactions between the two populations of classical and balanced cells are symmetric, i.e. each neuron sends and receives local excitation and long-range inhibition. **c-d**, Simulated HD tuning curves at different angular velocities for classical (c) or balanced (d) cells residing on different rings (blue – central rings, green – left rings, orange – right rings). The central column of plots corresponds to absence of head turns. The left plots correspond to increasing counterclockwise velocity, whereas the right plots correspond to increasing clockwise velocities. Note symmetric and asymmetric modulations of HD tuning as a function of angular velocity. **e-f**, Angular-velocity tuning of classical (e) or balanced (f) cells. Classical cells display strong angular-velocity tuning (gain of spikes), whereas balanced cells did not show angular-velocity tuning even though their HD tuning scaled with angular velocity (redistribution of spikes). Note that in the data there were more redistribution cells than gain cells, and in the model we took this into account by appropriately rescaling the synaptic weights of classical and balanced populations (Methods).

In our 6-ring model, classical cells showed HD-tuning scaling by angular velocity via addition of spikes (Fig. 5c,e), as we also observed in the classical 3-ring model (Fig. 1b-c). Importantly, balanced cells in the 6-ring model showed HD-tuning scaling via redistribution of spikes (Fig. 5d,f, note absence of AV-tuning in f). In close correspondence with the experimental results, we found that both classical and balanced neurons in the 6-ring model showed either symmetric scaling of their HD tuning for fast turns in both directions, or asymmetric scaling for clockwise/counterclockwise turns (Fig. 5c-d – compare orange, green, and blue curves). Notably, neurons with symmetric scaling for both clockwise and counterclockwise turning-directions were observed on the central ring only (Fig. 5c-d, blue). By contrast, neurons with asymmetrical scaling for either clockwise or counterclockwise turning-directions were observed exclusively on the right or left rings, respectively (Fig. 5c-d, green and orange). This suggests that HD cells with different shapes of HD-tuning scaling might be part of different subnetworks corresponding to different rings. Taken together, we showed that a novel recurrent 6-ring network composed of both classical and balanced neurons can account for the various forms of HD-tuning scaling that we observed in the bat dorsal presubiculum.

Finally, we explored what advantages could these additional populations of balanced cells bring to the overall function of the HD circuit. Directional sensory cues can update the position of the bump and correct the directional estimate that would otherwise accumulate errors over time^8, 28^. Therefore, the HD system would benefit from being sensitive to the appearance of directional sensory cues, even when such cues are weak (e.g. a dim and localized visual cue in a dark environment). We found that when the circuit is composed of classical cells only (as in the 3-ring model), a localized weak sensory-input is not sufficient to update the position of the bump (Supplementary Fig. 8a). This happens because classical cells are strongly suppressed outside of their preferred direction, and therefore a weak sensory stimulus arriving far-away from the current bump position would remain subthreshold and fail to impact the bump^47^. The addition of balanced cells that operate above spiking-threshold (as in the 6-ring model) allows efficient update of the position of the bump in the presence of weak sensory input (Supplementary Fig. 8c), because these cells can fire outside of their preferred-direction. Notably, the sensory-input initially leads to a change in activity of balanced cells only, but the recurrent interactions in the 6-ring model then ensure a subsequent coherent change in the activity of the classical cells as well (Supplementary Fig. 8c). This demonstrates that combining classical (gain) and balanced (redistribution) cells can increase the overall sensitivity of the HD network to directional cues.

## Discussion

Here we found that the head-direction tuning of neurons in the bat dorsal presubiculum was dynamically controlled by angular velocity. In many neurons, which were completely untuned when measured in a conventional way, a strong directional tuning emerged only at a specific range of angular velocities. In addition, the HD tuning was typically scaled by angular velocity, yielding a more pronounced directional tuning at higher angular velocities. In many neurons (63.7% of HD cells), HD-tuning scaling did not result from a change in the overall number of spikes emitted by the neuron (gain of spikes) but from a redistribution of spikes – which maintained the net spike count of the neuron. We showed that classical ring models of HD cells could not explain HD-tuning scaling that involves redistribution of spikes. However, we found that HD-tuning scaling by both gain and redistribution of spikes was recapitulated by a new type of attractor network model for HD cells, which consists of distinct populations of neurons residing on 6 recurrently-connected rings. Using this model, we were able to recapitulate *all* the different forms of HD-tuning scaling by angular velocity that we found experimentally – including scaling via gain and via redistribution of spikes.

We found here that the large majority of cells recorded in the dorsal presubiculum (61.3%) carried a significant HD signal. However, for most neurons, the directional tuning emerged only at some range of angular velocities. In particular, many cells showed the sharpest HD tuning at faster angular-velocities – where the animal spends relatively little time. Therefore, the conventional analysis that computes the directional response over the entire recording session (thus averaging across all angular velocities) is likely missing the directional tuning of many cells. Indeed, the percentage of significant directionally-tuned cells that we found – 61.3% of all presubicular cells – was much higher than in previous studies of HD cells in rats and bats^2, 3, 40, 41, 43^.

Here we demonstrated that the majority of neurons in the dorsal presubiculum showed HD tuning modulation by angular velocity, while maintaining a fixed spike count as a function of angular velocity. In these *redistribution cells*, HD-tuning scaling by angular-velocity resulted from a redistribution of spikes at the null direction versus the preferred direction – without a net change in the total number of spikes emitted by the neuron as a function of angular velocity. The fact that these neurons appear to be un-tuned to angular-velocity, when only the marginal tuning is considered, might explain why no conjunctive tuning to head-direction×angular-velocity was found to date in rat dorsal presubiculum^40^. Moreover, to our knowledge, such conjunctive tuning via redistribution of spikes, which we found here, was not reported to date in mammalian neural circuits.

Existing ring-attractor models of HD tuning could not explain our finding of neurons with conjunctive head-direction×angular-velocity tuning via spike redistribution. To explain the existence of redistribution cells, we proposed a 6-ring model with additional neurons that receive strong external excitation and strong feedback inhibition, keeping the neurons in a suprathreshold input-output regime. While we modeled classical (gain) and balanced (redistribution) cells as distinct classes of HD cells, it is possible that these neurons represent some extremes of a continuum of operational regimes. Our modeling results demonstrated that the presence of redistribution cells operating in the suprathreshold regime increased the sensitivity of the HD system to weak sensory signals. Therefore, redistribution cells could play a role in error-correcting mechanisms aimed at updating the bump position based on sensory inputs.

Our model generates a testable prediction regarding membrane potential dynamics of classical and balanced cells (see explanation about the prediction in Supplementary Fig. 9). This would allow testing whether the experimentally-recorded gain and redistribution cells indeed exhibit the subthreshold dynamics that is predicted by our model for the classical and balanced cells, respectively. Specifically, we predict that classical (gain) cells with asymmetric (clockwise or counterclockwise) shapes of their HD-tuning scaling would exhibit velocity-dependent modulation in their mean membrane-potential – whereas balanced (redistribution) cells would not show a net change in their mean membrane-potential as a function of angular velocity (Supplementary Fig. 9, compare panels b and d). Future work involving identification of the functional-anatomical properties of HD cells in mammals^27, 39, 48^ would allow testing this prediction, and also test whether gain and redistribution cells map onto genetically distinct cell types.

All existing models of HD cells rely on angular-velocity modulation of HD tuning to support angular-velocity integration – a mechanism that allows updating the internal estimate of direction (bump of activity) according to head turns^10, 22–24, 26, 29–31^. This mechanism was recently supported by findings in the *Drosophila* HD system^35^. In the mammalian brain, only a small fraction of HD cells was reported to show such modulation by angular velocity. These neurons were found in the dorsal tegmental nuclei^12, 14^, leading to the view that the generation and update of the HD signal by self-motion cues happen subcoritcally^30^. Our finding of a prominent modulation of the HD-tuning by angular velocity in the dorsal presubiculum suggests that a parallel computation might occur also at the cortical level. Since the dorsal presubiculum receives inputs from visual areas, it might be involved in angular-velocity integration based on optic flow^9, 49, 50^.

Finally, what are the possible advantages of having neurons that exhibit angular-velocity modulation of the HD tuning – in addition to potentially supporting cortical angular-velocity integration? First, we have previously shown that angular-velocity modulation of bat 3D HD cells allows shifting the coding scheme towards conjunctive tuning of azimuth and pitch at faster angular velocity^42^ – in a manner that can optimize population coding according to animal behavior^42, 51, 52^. Second, unless the HD-tuning scales with angular velocity, HD cells would emit very few spikes during fast head-turns – because the head of the animal would sweep very rapidly through the preferred direction of the neuron. Therefore, the observed upscaling of the HD tuning at high angular-velocities could comprise a compensatory mechanism that maintains a roughly constant number of spikes emitted at the preferred direction, irrespective of angular velocity. Taken together, the dynamic control of HD tuning by angular velocity, which we found here, could support cortical angular-velocity integration – and provide a reliable representation of the HD signal across a wide range of angular velocities.

## Acknowledgments

We thank J. Aljadeff, R. Darshan, A. Rubin, B. Hulse, and L. Las for comments on the manuscript; B. Pasmantirer and G. Ankaoua for mechanical designs; S. Kaufman for bat training; and A. Tuval and M. Weinberg for veterinary care. This study was supported by research grants to N.U. from the European Research Council (ERC-StG – NEUROBAT and ERC-CoG NATURAL_BAT_NAV), the Israel Science Foundation (ISF 1319/13), and the Minerva Foundation; and by a Clore predoctoral excellence fellowship to A.F. The data and computer code are archived on the Weizmann Institute of Science servers and will be made available upon reasonable request.

## Methods

### Subjects and behavioral training

Neural activity was recorded from the dorsal presubiculum of 4 adult male Egyptian fruit bats, *Rousettus aegyptiacus*. All experimental procedures were approved by the Institutional Animal Care and Use Committee of the Weizmann Institute of Science, and are detailed elsewhere^41^. Briefly, bats were trained to crawl in search of randomly distributed food pieces on the floor of a square wooden box (50 × 50 cm). The experiment included 2 crawling sessions, each lasting 15–25 min, conducted ~1 hour apart from each other, and were flanked by two sleep sessions, lasting ~10 min each. The data from these 4 bats were previously published in ref.^41^.

### Surgery, recording techniques, spike sorting, and video tracking

All surgical and recording procedures were as described previously^41^. Briefly, after completion of training, bats were implanted with a four-tetrode microdrive (4-drive, Neuralynx; weight 2.1 gr). While the bat was under isoflurane anesthesia, a circular opening (craniotomy diameter of 1.8-mm) was made in the skull over the right hemisphere. The center of craniotomy was positioned over the dorsal presubiculum, 3.3 mm lateral to the midline and 3.15 mm anterior to the transverse sinus that runs between the posterior part of the cortex and the cerebellum (R. Eilam, L. Las, M. P. Witter, and N. Ulanovsky, *Stereotaxic brain atlas of the Egyptian fruit bat*, in preparation).

Following surgery, tetrodes were slowly lowered towards the dorsal presubiculum. The tetrodes were always moved at the end of each recording day (40–200 μm daily), in order to obtain recordings from new ensembles of neurons daily. For each bat, one tetrode was left in an electrically-quiet zone and served as a reference, and the remaining three tetrodes served as recording probes. During recordings, a unity-gain preamplifier (HS-18, Neuralynx) was attached to a connector on the microdrive. Signals were amplified (×1,400–5,000) and band-pass filtered (600–6,000 Hz, Lynx-8 or DigitalLynx, Neuralynx), and a voltage-threshold was used for collecting 1-ms spike waveforms, sampled at 32 kHz. Data were collected continuously throughout all the behavioral and sleep sessions of each recording day (1.5–2 hours daily).

Spike-sorting procedure was described previously^41^. Briefly, spike waveforms were sorted on the basis of their relative energies and amplitudes on different channels of each tetrode (SpikeSort3D, Neuralynx). Data from all sessions, including the two sleep sessions, were spike-sorted together. Well-isolated clusters of spikes were manually encircled (‘cluster-cutting’), and a refractory period (<2 ms) in the interspike-interval histogram was verified.

Video-tracking was as described previously^41^. Briefly, a single camera that was placed vertically overhead the arena, was connected to a video-tracker (Neuralynx) with a 25-Hz sampling rate, which was synchronized to the neural recordings. The video tracker recorded the positions of an array of light-emitting diodes (LEDs) of different colors positioned on the bat’s head. The pair of diodes positioned along the antero-posterior axis of the head was used to estimate the animal’s head direction in the azimuthal plane (direction of the nose), which was the plane used for all analyses of HD in the current work.

### Angular-velocity computation and binning

*Angular velocity* was computed as the time derivative of the azimuthal HD angle, by taking the difference in the azimuthal HD angle between each consecutive video frame and multiplying it by the frame rate. To examine the effects of angular velocity on the mean firing rate and on the HD tuning of the neurons, in all the analyses we binned the data into angular-velocity bins using a sliding window of 25°/s, which we moved in steps of 1°/s (e.g. in Fig. 2a-iii,iv). We analyzed angular-velocity bins whose centers spanned the range of [−50°/s, +50°/s], with positive and negative angular velocities corresponding to clockwise and counterclockwise head-turns, respectively. In some cases, bats turned their head at slightly higher angular velocities, but these epochs were typically too brief to be used for constructing a full HD-tuning curve. Within the analyzed range of [−50°/s, +50°/s], we considered a specific angular-velocity bin as valid if it contained at least 30 seconds of behavioral data. Non-valid bins were discarded from the analysis.

### Inclusion criteria

A total of 266 well-isolated neurons were recorded from dorsal presubiculum of 4 bats during 2 crawling sessions. Of these 266 neurons, we included for analysis a total of 346 cells×sessions, based on the following criteria: (i) The cell emitted at least 150 spikes in the relevant behavioral session. (ii) The relevant behavioral session included valid angular-velocity bins whose centers spanned at least a range of [−25°/s, +25°/s]. Note that most sessions that were included in the analysis, had enough behavioral data over the entire range of [−50°/s, +50°/s] (see examples in Supplementary Fig. 2a-b, column v). Throughout the paper, we referred to cells×sessions as cells. These inclusion criteria were applied to all neurons – including conventional HD cells, angular-velocity tuned cells, and cells with HD tuning at some angular velocity (green, red and blue circles in the Venn diagram in Fig. 3c, respectively).

### Computing conventional HD-tuning curves over the entire session

We first computed the *conventional HD tuning* (Fig. 2a, column *i*) of a neuron over the entire session, irrespective of angular velocity, as follows. We binned the 360° of the azimuthal HD angles into 18 bins of 20° each, and computed the firing rate in each bin by dividing the number of spikes in that bin by the time the animal spent in that bin. The resulting tuning curve was smoothed in a circular manner using a rectangular window of size 5 (the smoothing was done on the firing-rate curve, i.e. after dividing the unsmoothed spike-curve by the unsmoothed time-spent curve). Unvisited bins (where total time-spent, before smoothing, was < 0.5 sec) were discarded from the analysis.

The directionality of the tuning curve was quantified by computing the Rayleigh vector length of the circular distribution, using the following definition^53^:

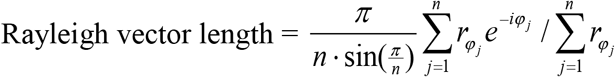

Where *n* is the number of circular HD bins, φ_*j*_ is the direction in radians of the *j*-th circular bin (namely, 2 *πj* / *n*), and *r_φ_j__* is the average firing rate given the animal’s HD.

In order to identify neurons with significant conventional HD tuning we applied a shuffling-based significance test on the Rayleigh vector length^4, 5, 41^. Significance was determined for each neuron separately: Specifically, for each recorded neuron, the entire sequence of spikes was time-shifted by a random (uniformly-distributed) time interval, with the end of the session wrapped to the beginning. The minimal time interval for the shift was 10 s, and the shift was no longer than the total session duration minus 10 s. This preserved the spike number and the temporal structure of the neuron’s firing pattern, but dissociated the time of spiking from the animal’s actual behavior. This shuffling procedure was repeated 1,000 times for each neuron. A session was defined as having a conventional HD tuning if its Rayleigh vector length exceeded the 95^th^ percentile of the shuffled distribution for this session, and was larger than 0.15. The total number of cells×sessions with conventional HD tuning is depicted in Fig. 3c (green circle).

### Evaluating the HD tuning at different angular velocities

To analyze HD tuning at different angular velocities (Fig. 2a, column *ii*), we divided the data into angular-velocity bins, as described above. We then computed the HD-tuning curve for each angular-velocity bin. HD-tuning curve for each angular-velocity bin was computed in the same way as for the ‘conventional tuning’, except that now we used only spikes and behavioral data that corresponded to that particular angular-velocity bin.

To assess HD in each angular-velocity bin, we first fitted HD-tuning curve with a circular normal function, also known as the von Mises function – which has the following expression:

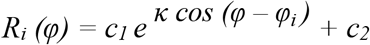

Where *φ* is the preferred direction of this neuron in radians, *φ_i_* is the direction in radians of the *i*-th circular bin, and *κ*, *c*_1_, and *c*_2_ are constants.

Using the von Mises fit, we measured the following parameters of the HD-tuning curve: *HD-tuning quality* (Fig. 2a, column *iii*) was defined as the goodness-of-fit of the HD-tuning curve in each angular-velocity bin to the corresponding von Mises function. Based on the goodness of fit we divided the bins into: ‘tuned’ (*R*^2^ ≥ 0.75) or ‘untuned’ (*R*^2^ < 0.75).

*Preferred direction* (PD) was defined as the angle corresponding to the peak of the von Mises function fitted to the HD-tuning curve (Supplementary Fig. 1a).

*Preferred-direction firing rate* was defined as the maximal firing rate of the fitted HD-tuning curve.

*Null-direction firing rate* was defined as the minimal firing rate of the fitted HD-tuning curve.

*Modulation depth* was defined as the difference between the preferred-direction firing rate and null-direction firing rate (Supplementary Fig. 1a).

*HD-tuning scaling* was defined as a change in the modulation depth of the tuning as a function of angular velocity (see schematic illustration in Supplementary Fig. 1 and examples in Fig. 3a-b, column iii). It consisted of modulation-depth values at each angular-velocity bin versus the angular velocity in these bins. The resulting curve was smoothed using a rectangular window of size 12°/s.

To analyze changes in the preferred direction of a neuron (Fig. 2a,d), we computed the circular standard deviation^54^ of the preferred directions of each cell at different angular-velocity bins. This computation was done separately for angular-velocity bins for which each cell was tuned (Fig. 2d, magenta) or untuned (black), according to the tuning quality criterion above. For evaluating the stability of the preferred direction across the angular-velocity range (Fig. 2e), we computed the mean preferred direction of all tuned angular-velocity bins, separately for bins corresponding to clockwise and counterclockwise turning-directions, and took the difference between the two means. For this analysis, we included only cells that were tuned over an angular-velocity range that spanned at least ±15 °/s (92 out of 212 cells×sessions with significant HD tuning at some angular-velocity bin).

### HD-tuning significance in a particular angular-velocity bin

To assess the significance of directional tuning in a particular angular-velocity bin, we performed a shuffling procedure on the Rayleigh vector length for that bin. For the shuffling for each angular-velocity bin, we used only spikes and behavioral data from that particular angular-velocity bin. Note that the behavioral data in each angular-velocity bin were intermittent in time, because they were composed of different time-epochs during the session, in which the animal’s head was moving at that specific angular-velocity range. Therefore, for this shuffling procedure, we concatenated all the relevant epochs of the behavioral data and spikes that corresponded to that bin. We then shifted the times of all spikes for that bin rigidly by a random time interval (uniformly-distributed, with the end wrapped to the beginning); the time-shift was at least 5 s, and no more than the total duration available for that bin minus 5 s. The shuffling was repeated 1,000 times for each angular-velocity bin. In all other aspects, the shuffling procedure was the same as the shuffling for the conventional HD tuning.

We considered a cell×session to have a significant HD tuning at some angular velocity if it matched all of the following criteria (a total of 212 cells × sessions [61.3% of all recorded neurons] matched these criteria): (i) The Rayleigh vector length of the HD tuning in at least one angular-velocity bin exceeded the 95^th^ percentile of the shuffled distribution for that bin in this session (we note that in practice, almost all neurons that matched all the criteria [195/212 cells×sessions] had significant HD tuning in ≥5 angular-velocity bins). (ii) The Rayleigh vector length at that same angular-velocity bin was > 0.15. (iii) The HD tuning curve at that bin was stable. Stability was determined by computing the Pearson correlation coefficient between HD tuning for even versus odd quartiles of the data, and we required a Pearson correlation of: *r* > 0.5. Note that most of the neurons that matched all the above criteria, were classified as non-HD cells by the conventional HD analysis (i.e. when the tuning curve was computed over the entire session).

### Assessing the effect of random fluctuation on HD-tuning scaling at different angular velocities

To assess whether the observed HD-tuning scaling by angular velocity could be an artifact of behavioral sampling biases or random fluctuations in the firing rate, we modeled the HD firing in each angular-velocity bin assuming a conventional HD tuning with Poisson noise. To this end, for each angular-velocity bin, we first divided the behavioral data into 40-ms time intervals (corresponding to the 25-Hz frame-rate of the video tracker), and created a vector of HD angles in each 40-ms time interval based on the session’s behavioral data. We next computed the expected number of spikes during each time interval, by taking the conventional HD tuning curve (computed for the entire session) as the rate-parameter for a Poisson process. We used these synthetically-generated spikes and real behavioral data to compute a new HD-tuning curve for each angular-velocity bin, as described in the previous section. For each angular-velocity bin, we repeated this process to generate 100 HD-tuning curves, and used them to compute the distribution of modulation-depth values expected by chance at each angular-velocity bin.

We used ±1 standard deviations around the mean of the modulation-depth distribution (in each angular-velocity bin) to define a confidence interval for HD-tuning scaling, assuming a conventional HD tuning with no genuine modulation by angular velocity (Supplementary Fig. 2a-b, column iii – see shaded gray area). Scaling of HD tuning that occurred within the confidence interval (within the shaded gray area) could be fully explained by random fluctuations of the firing rate of the conventional HD-tuning curve – and therefore we considered such modulations as non-genuine changes in the HD tuning (Supplementary Fig. 2b, column iii – note that in these examples of non-genuine modulations, HD-tuning scaling (blue line) was within the shaded gray area for almost all angular velocities). By contrast, angular-velocity bins with HD-tuning scale values outside of the confidence interval were considered genuinely scaled by angular-velocity (Supplementary Fig. 2a, column iii – note the substantial ranges of angular velocity where the blue line was outside of the shaded gray area).

A session was considered to exhibit a significant HD-tuning scaling by angular velocity, if: (i) At least 10 of the bins of the scaling modulation curve were outside of the confidence interval (Supplementary Fig. 2d: neurons to the right of the gray line). (ii) The maximum modulation-depth value was at least 75% above the minimal modulation-depth value (Supplementary Fig. 2a, column iii). Curves with modulation < 75%, were not considered to be modulated. Example neurons without a significant HD-tuning scaling are shown in Supplementary Fig. 2b (column iii).

### Angular-velocity tuning

Angular-velocity tuning of a neuron was defined as the relation between the firing-rate in each angular-velocity bin and the angular velocity, without taking HD-tuning in this bin into account. Firing rate in each angular-velocity bin was computed by dividing the number of spikes in that angular-velocity bin by the total duration that bat spent in that angular-velocity bin. The resulting AV-tuning curve was then smoothed using a rectangular window of size 12°/s.

To assess the significance of AV tuning, we performed a shuffling procedure using all spikes, as described above in the section on the conventional HD tuning. For each shuffle realization (1,000 shuffles in total) we computed the firing rate in each angular-velocity bin and thus constructed the distribution of the expected firing-rate values at each angular-velocity bin, assuming that there is no genuine AV tuning. We used the firing rate ± 1 standard deviations above or below the mean (separately for each angular-velocity bin) to define a confidence interval for changes in the firing rate as a function of angular-velocity.

A cell×session was considered to exhibit a significant AV tuning (i.e. a significant modulation of its firing rate by angular velocity), if: (i) At least 10 bins of the AV-tuning curve were outside of the confidence interval. (ii) The maximum modulation value was at least 75% above the minimal modulation value. Curves with modulation below 75% increase, were considered to be untuned.

### Classification of HD cells according to the shape of their modulation by angular velocity

We considered for this classification only neurons that had a significant HD tuning in at least some angular velocity (61.3% of all recorded neurons), as follows. For each cell, we computed the Pearson correlation coefficient between the shape of HD-tuning scaling and one of three templates depicted in Fig. 4a (e.g. symmetric scaling, column 1 and row 1; asymmetric scaling, columns 2-3 and rows 2-3). A similar calculation of the Pearson correlation coefficient was done for the shape of AV-tuning. Classification was determined according to the template that yielded the highest value of the Pearson correlation coefficient.

We considered a session as belonging to one of these classes, if: (i) The Pearson correlation coefficient with the best template was *r* > 0.5. (ii) There was a minimal number of bins outside the confidence interval, as described above. (iii) The maximum modulation was greater than 75%. If the Pearson correlation with all of the three templates was lower than 0.5 (or if there were not enough angular-velocity bins outside of the confidence-interval), yet the maximum modulation exceeded 75%, then the neuron was classified as having ‘other’ modulation shape (Fig. 4a, column 5 ‘other’ and row 5 ‘other’). When the neuron’s tuning was not classified according to one of these four categories, the neuron was considered to have no modulation by angular velocity (represented by a flat line icon in Fig. 4a, see column 4 and row 4).

### Gain and Redistribution cells

Neurons were defined as *gain cells* if the shapes of their HD-tuning scaling and AV-tuning belong to the same class. Specifically, gain cells were neurons in the first three diagonal bins in Figure 4a (i.e. with symmetric scaling, [column 1, row 1]; and asymmetric scaling [columns 2-3, rows 2-3]). Neurons were defined as *redistribution cells* if they had a significant HD-tuning scaling but no AV-tuning. Specifically, redistributions cells were neurons in the 4^th^ row in Figure 4a (row 4, columns 1-3,5).

### Assessing stability across time of HD cell-activity modulation by angular velocity

For stability analysis (Supplementary Fig. 4), we divided each angular-velocity bin into four time-epochs. We constructed HD-tuning scaling separately for odd versus even data-epochs, and determined stability of HD-tuning scaling modulation by computing the Pearson correlation coefficient between the two. We did analogous computations to assess the stability of AV tuning. For the analysis of the stability of HD-tuning scaling, we used only angular-velocity bins that were outside of the confidence interval for significant HD-tuning scaling computed for the entire epoch (Supplementary Fig. 2, iii - where the confidence interval was computed as described above), and contained at least 15 seconds of data for odd epochs and at least 15 seconds for even epochs. This was done to avoid comparing the stability of HD-tuning scaling for angular-velocity bins that were governed by noise fluctuations or had insufficient sampling.

## Methods for the six rings angular integration circuit with scaling and redistribution phenomena

We consider networks of neurons that are connected with the symmetry of a ring. We first analyse a simplified model that can account for the gain and redistribution tunings. We then describe a more complex model that can perform angular integration.

### 1 Classical and balanced ring networks

We consider two populations of neurons described by their firing rates at time *t, f_g_*(*θ, t*) and *f_r_*(*θ, t*) corresponding respectively to the “classical” (gain) and “balanced” (redistribution) neural populations. Each neuron is associated with an angle *θ* between 0 and 2π along the ring. We will describe the dynamics with continuous angular variables for simplicity. In the simulations, the angles are discretized with a grid of 64 uniformly sampled angles. We use, for all neurons, a threshold-linear relationship between the input current *I* to a neuron and its firing rate *f, f* = [*I*]_+_.

We propose that the average firing rate of the redistribution population is imposed by the balance between a strong recurrent inhibitory input and a strong external input. Without additional inputs and interactions, the dynamics for the classical (gain) population *f_g_* and the balanced (redistribution) population *f_r_* obey the following equations at the leading order in the large parameter *γ*:

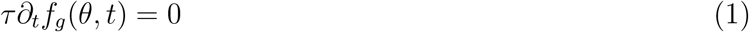

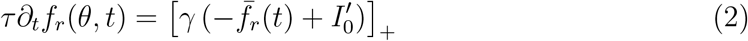

where *γ* ≫ 1 (*γ* = 100 unless otherwise specified), 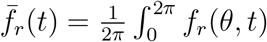 is the average activity across the balanced neurons, and τ is the time constant of neuronal integration. Hence, the mean activity of the redistribution population is clamped to 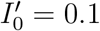.

The localized activity on the ring arises from localized excitation and long-range inhibition. To preserve the rotation symmetry along the ring, the interaction between neurons is expressed as a convolution product, with a kernel chosen to be a Von Mises function. The dynamical equations for the classical and balanced populations are then expressed as follows at the first order in *γ*:

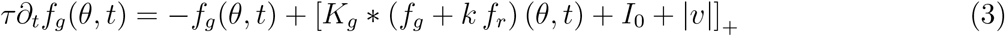

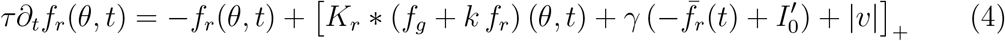

The convolution between *K* and the firing rate of the neurons *f* is defined as:

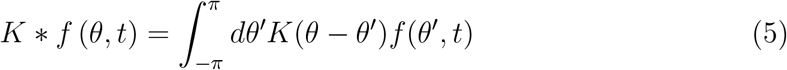

Hence, *K*(*θ* – *θ*′) is the synaptic weight between the neurons with associated angles *θ* and *θ*′. We chose a convolution kernel of the form *K*(*θ*) = *J*_0_ + *J*_1_*p*(*θ*; *κ*), with *p*(*θ*; *κ*) the Von Mises distribution of parameter *κ* (*κ* = 5 unless otherwise specified):

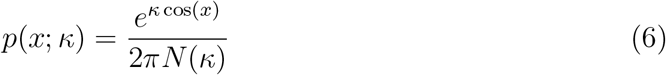

where *N*(*κ*) is the modified Bessel function of order 0. The parameters *J*_0_ and *J*_1_ in the convolution kernels describe respectively the strength of the global inhibitory interaction and of the head-direction specific interaction. The intensity of the input is chosen as follows:

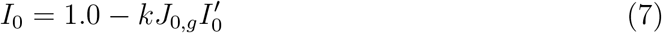

Redistribution cells were more numerous compared to gain cells in the data. The factor *k* accounts for the ratio between redistribution and gain cells. In our simulations, we used *k* = 3, however the results of our simulations do not critically depend on the specific value of this parameter.

Both populations receive an additional input that depends on the angular velocity *v*. In order for the balanced population to keep a constant mean firing rate, this velocity input is chosen to be of order 1.

Supplementary Figure 6 shows tuning curves for the classical and balanced populations for a range of input angular velocities |*v*|.

The parameters chosen for this numerical simulation are as follows:

|  | Classical (gain) | Balanced (redistribution) |
| --- | --- | --- |
| $J_0$ | -0.7 | 0.0 |
| $J_1$ | 0.7 | 0.25 |

### 2 Angular velocity integration circuits

Our model (Figure 5a) consists of several populations of neurons representing the azimuth of the bat head direction. We first consider a model of classical (gain) c ells. This model consists of a central ring of neurons and two side rings. The model exhibits localized persistent activity and performs angular velocity integration upon the activation of the side rings by an angular velocity signal. We next consider an extension of this model that includes the balanced (redistribution) cells.

#### 2.1 Model of angular integration with three classical populations

We assume that the neurons of the central ring (population *C*) establish excitatory synapses with the neurons of the side rings (populations left *L* and right *R*) representing similar head direction. The side ring cells in turn excite, with an offset ±Δ, the central ring. Moreover, neurons of the central ring inhibit all the neurons of the side rings with identical strength (global inhibition).

As before, neurons are indexed by an angle *θ* between 0 and 2*π* along each of the three rings, with neural activities at time *t* described by the firing rates *f_C_*(*θ, t*), *f_L_*(*θ, t*) and *f_R_*(*θ, t*) for the central, left, and right rings respectively, with dynamics:

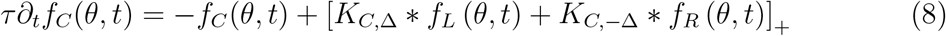

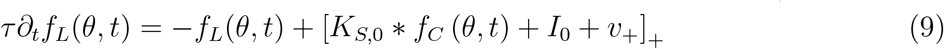

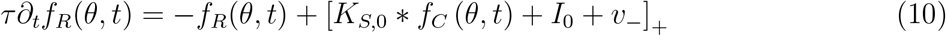

As before, since the model is rotation symmetric, the coupling between the different populations is expressed as a convolution product. The kernels are expressed as follows:

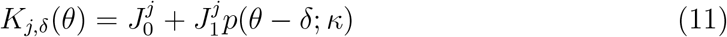

where *p*(*θ, κ*) is the Von Mises distribution (Eq. 6), the index *j* specifies the population of neurons that receives the input: center *C*, or side *S* rings (either left or right), and *δ* denotes the offset of the interaction (*δ* = 0, −Δ, +Δ).

The constant intensity input is chosen as I_0_ = 1.0. The angular velocity inputs on the side rings are uniform among the populations and have the following form as a function of the external velocity *v*:

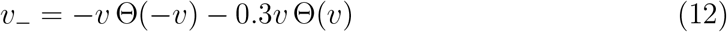

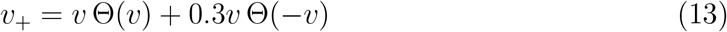

where Θ(*x*) is the Heaviside function (Θ(*x*) = 0 for *x* < 0 and Θ(*x*) = 1 otherwise).

The parameters that are used in the simulation presented on Figure 1 are as follows:

|  | Central to side | Side to central |
| --- | --- | --- |
| $J_0$ | -1.5 | -0.4 |
| $J_1$ | 2.5 | 2.5 |

and the angular shift was Δ = *π*/8.

#### 2.2 Six populations model with classical and balanced cells

The complete model combines the classical and balanced populations with the previous circuit performing the angular velocity integration. Thus for each population of the previous simulation, we add another population that receives a strong external input and strong recurrent inhibition, similarly to the balanced population of Section 1. The new system is described by the dynamics:

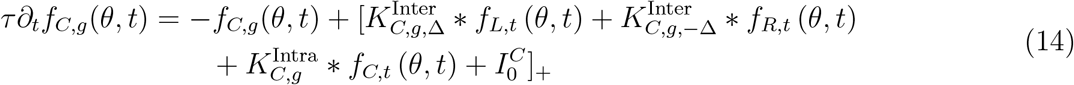

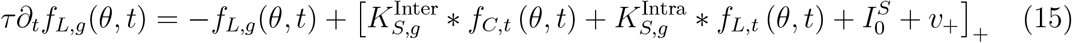

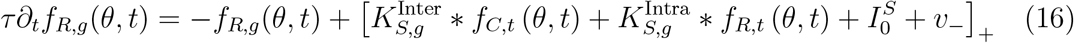

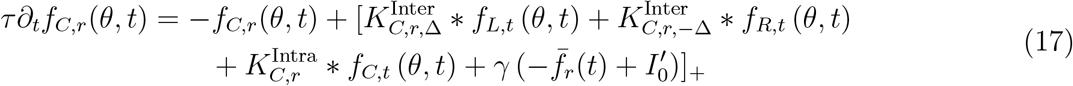

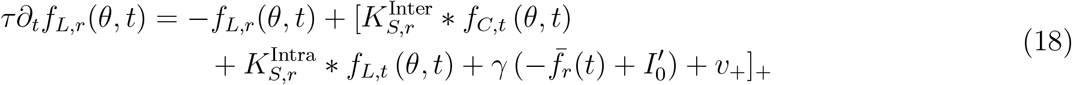

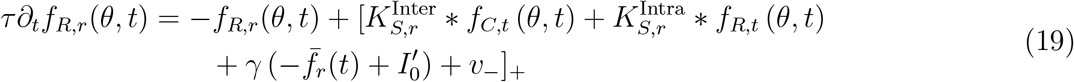

In these equations, firing rates are denoted by *f_α,β_*(*θ,t*), where *α* indexes the central (*C*), left (*L*), or right (*R*) rings, while *β* indexes classical (*g*), balanced (*r*), or both (*t*) populations. The combined firing rate from both classical and balanced populations is defined as *f_α,t_* = *f_α,g_* + *kf_α,r_*, where *k* is the ratio between the number of classical and balanced cells.

The convolution kernels describing the interaction between neurons are denoted as:

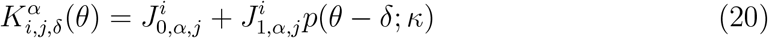

where *p*(*θ,κ*) is the Von Mises distribution (Eq. 6), the index *i* specifies the ring of neurons that receives the input (center *C*, side *S*), *j* specifies the classical (*g*) or balanced (*r*) population, *δ* denotes the offset of the interaction (*δ* = 0, −Δ, +Δ), and *α* denotes within (*Intra*) or between (*Inter*) rings interactions.

The constant intensity inputs are chosen as 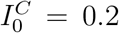 and 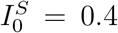. The input velocities on the side rings are uniform among the populations and have the following form as a function of the external velocity *v*:

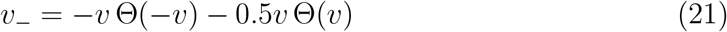

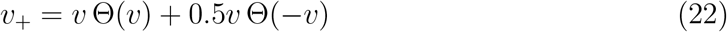

where Θ(*x*) is the Heaviside function (Θ(*x*) = 0 for *x* < 0 and Θ(*x*) = 1 otherwise).

The parameters that are used in the simulation presented on Figure 5 are as follows:

|  | Intra-connectivity |  | Inter-connectivity |  |
| --- | --- | --- | --- | --- |
|  | Classical | Balanced | Classical | Balanced |
| $J_0^C$ | -0.4 | 0.0 | -0.25 | 0.0 |
| $J_1^C$ | 0.3 | 0.05 | 0.75 | 0.05 |
| $J_0^S$ | -0.15 | 0.0 | 0.0 | 0.0 |
| $J_1^S$ | 0.55 | 0.05 | 0.75 | 0.05 |

and the angular shift was Δ = *π*/6.

For the simulation results reported in Supplementary Figure 8b, the population of classical neurons was removed from the network. In that setting, its topology actually correspond to the three-ring network of Section 2.1, with the addition of the strong external excitation and strong recurrent inhibition (the terms including *γ*). To generate HD tuning in this network, we increased the strength of this interaction as follows:

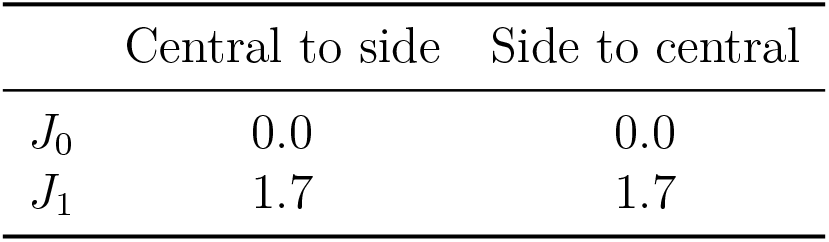

|  | Central to side | Side to central |
| --- | --- | --- |
| $J_0$ | 0.0 | 0.0 |
| $J_1$ | 1.7 | 1.7 |

and the angular shift was Δ = *π*/8.

### 3 Simulations including noise

To quantify the diffusion properties of the bump position with respect to the intrinsic noise, we modified the simulations as f ollows. For each firing rate variable (*C*(*θ_i_, t*) for instance) we associated a noisy variable 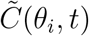 that followed a stochastic differential equation. These variables were used in the currents driving the recurrent dynamics (Eqs. 14–19). These dynamics effectively add a Poisson-like statistics (variance proportional to mean at steady state) to the firing rate:

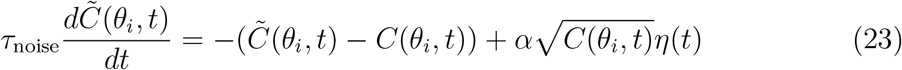

where *η*(*t*) is a gaussian white noise of variance 1: 〈*η*(*t*)〉 = 0 and 〈*η*(*t*)*η*(*t*′)〉 = *δ*(*t* − *t*′). *τ*_noise_ = 0.1*τ* and *α* = 0.0075 in the presented numerical simulations.

Simulations were performed with a modified stochastic Runge-Kutta algorithm.

### 4 Angular position resetting by weak, localized, external inputs

The bump position is reset with a static tuned input on the central balanced ring. The input is localized in *θ* according to a Von Mises function with *κ* = 20 and a prefactor of 0.15.

### 5 Software

All simulations and plots were performed with Python and its scientific packages `numpy, scipy` and `matplotlib`, and all codes are available on https://github.com/hrouault.

**Supplementary Fig. 1.**
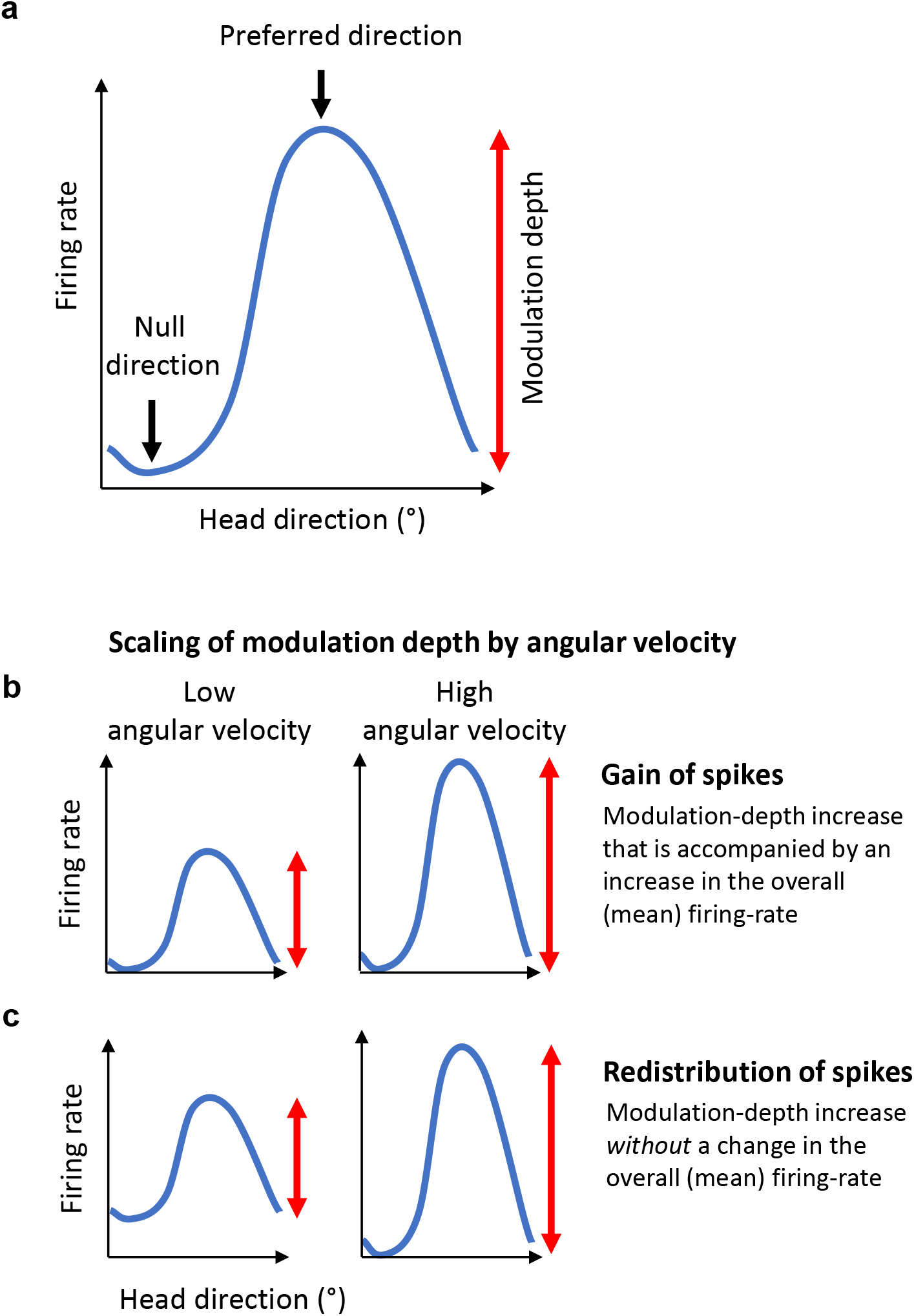
Schematic illustration of HD-tuning scaling by angular velocity via gain of spikes or via redistribution of spikes. **a**, Cartoon of a head-direction tuning curve, showing the definitions of the preferred direction (direction with maximal firing-rate), null direction (direction with minimal firing-rate), and modulation-depth (difference in firing-rates between the preferred and null directions). **b-c**, Cartoons of two types of HD-tuning scaling (rows) as a function of angular-velocity (columns): gain of spikes (b) and redistribution of spikes (c). **b**, Gain of spikes: an increase in modulation depth accompanied by an increase in total spike-count. **c**, Redistribution of spikes: an increase in modulation depth without any change in spike-count. For gain cells there is in an overall increase in mean firing-rate as a function of angular velocity (panel b, compare left versus right). For redistribution cells there is no change in mean firing-rate as a function of angular velocity – which is achieved because the increase in the firing-rate at the preferred-direction is accompanied by a corresponding decrease in firing-rate at the null-direction (panel c, compare left versus right).

**Supplementary Fig. 2.**
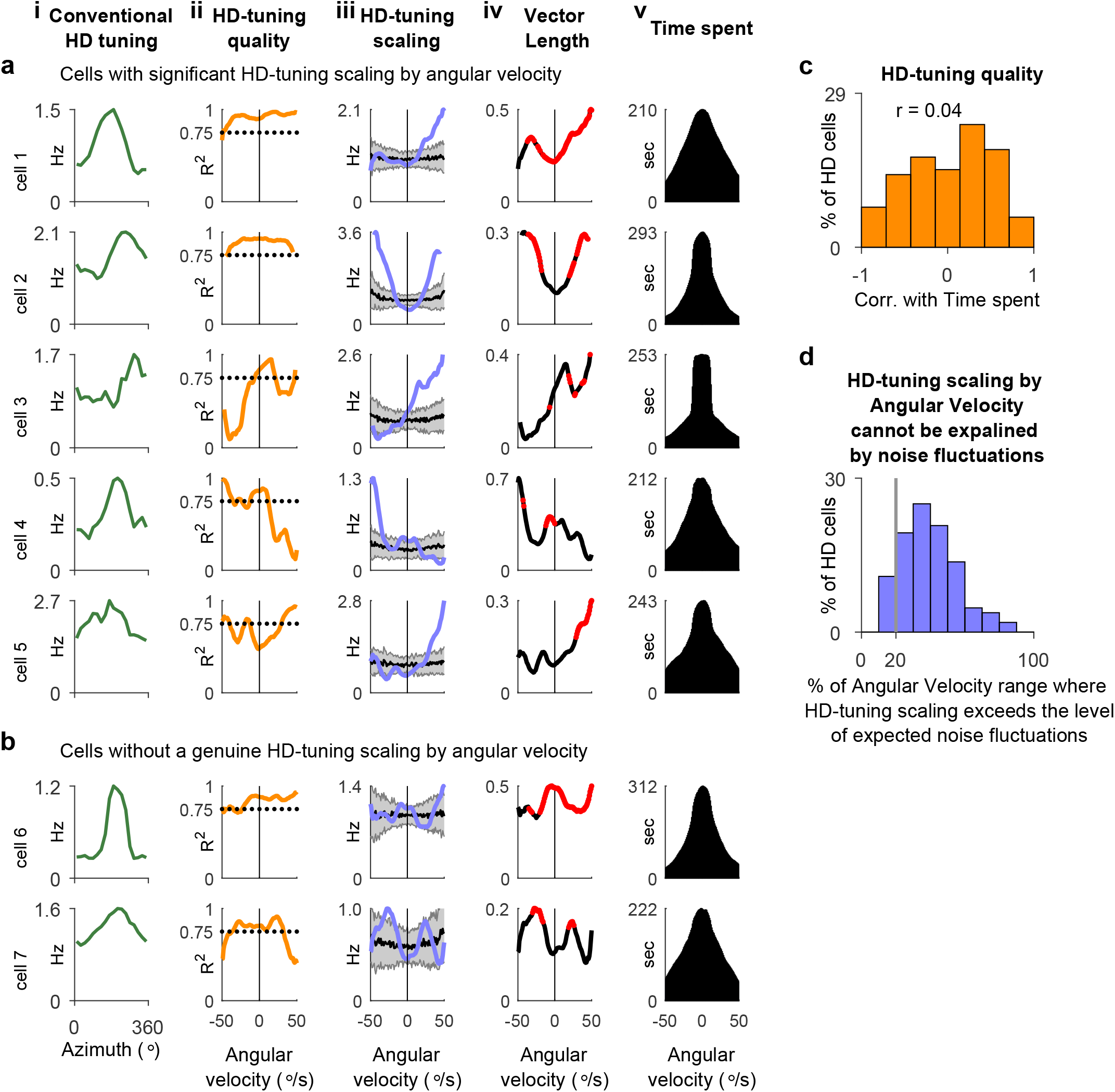
Determining the significance of head-direction tuning modulation as a function of angular velocity. **a-b**, Example cells (rows) with significant (a) or non-significant (b) modulations of head-direction (HD) tuning by angular velocity. Columns: *i*, Conventional HD-tuning curve, computed for the entire session. The parameters in columns ii-v, are plotted as a function of angular velocity: *ii*, Quality of HD tuning (assessed by goodness-of-fit to von Mises tuning); values with *R*^2^<0.75 were considered untuned. *iii*, Observed HD-tuning scaling (blue), overlaid with the expected variability in scaling (gray, mean ± s.d.) – where the expected variability was computed assuming Poisson firing-rate fluctuations (rather than genuine modulations; Methods). *iv*, Rayleigh vector length of the HD tuning. Red, angular-velocity bins in which the Rayleigh vector length significantly exceeded the value expected from shuffled distribution (Methods). *v*, Time spent by the bat in each angular-velocity bin that was used for analysis. Note in column *iii* that for neurons, which exhibited significant modulation of the HD tuning by angular-velocity (a), the scaling profiles of these neurons exceeded the confidence interval for random firing-rate fluctuations (blue curve emerged outside the gray shaded area in column iii). By contrast, for neurons without a genuine modulation of HD tuning by angular velocity (b), the apparent scaling could be entirely explained by random fluctuations in the firing rate (column *iii*: blue curve is inside the shaded gray area). **c**, Distribution of Pearson’s correlation coefficients between the time-spent at different angular velocities and the quality of HD tuning in these angular velocities. The distribution is around 0 (mean Pearson correlation: *r* = 0.04.), indicating that changes in tuning quality did not reflect insufficient time-spent at a particular angular velocity. **d**, Percentage of angular-velocity range where HD-tuning scaling by angular velocity exceeded the expected variability due to random fluctuations in firing-rate (see a-b, column *iii*).

**Supplementary Fig. 3.**
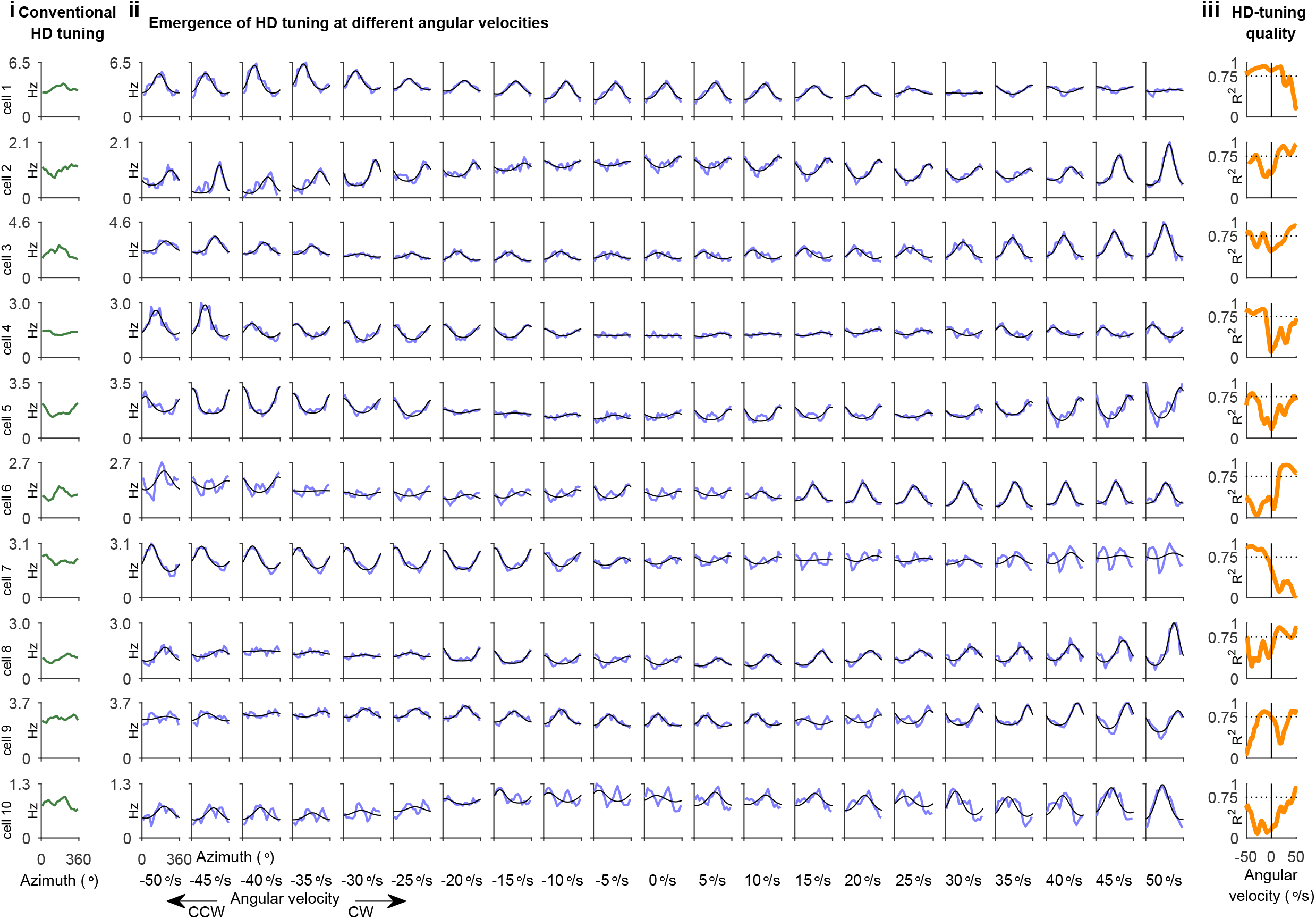
Head-direction tuning emerged at certain angular velocities in cells without a conventional head-direction tuning. Example cells (rows) showing the emergence of HD-tuning at different angular velocities (AV). Note that these cells did not have a significant conventional HD-tuning (computed for the entire session across all angular velocities: green curves in column i). Columns: *i*, Conventional HD tuning computed for the entire session. *ii*, HD-tuning curves (blue) computed for different angular-velocity bins, overlaid with von Mises fits (black). Numbers below each column indicate the center of the angular-velocity bin, plotted here in small jumps of 5 °/s. CW, clockwise; CCW, counterclockwise rotations of the head. *iii*, Quality of HD tuning (assessed by goodness-of-fit to the von Mises tuning), as a function of angular velocity. Note that in *ii*, there is an overlap between nearby columns, because the computation of HD curves was done using our standard angular-velocity binning of 25°/s (Methods); in all other figures throughout the paper, there is no such overlap, as columns are separated by 25°/s, and thus HD-tuning curves in different angular-velocity columns represent independent data.

**Supplementary Fig. 4.**
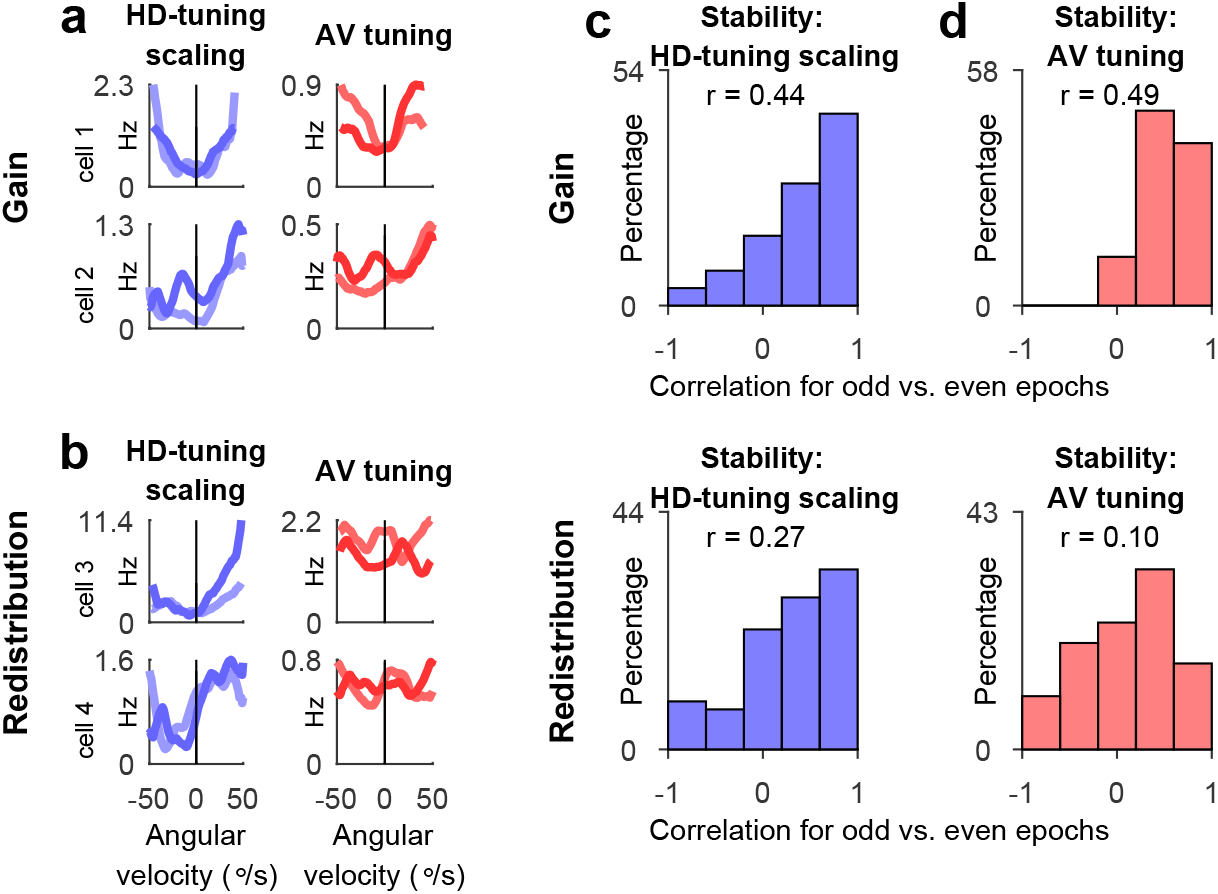
Stability across time of HD-cell activity modulation by angular velocity. **a-b**, Stability of HD-cell activity modulation by angular velocity, computed for odd versus even quartiles of a recording session (Methods), for two example gain cells (a) and two example redistribution cells (b). Overlaid curves represent HD-tuning scaling (left) and AV-tuning (right), computed for two separate parts of the behavioral sessions (odd versus even quartiles). **c**, Population stability analysis of HD-tuning scaling for gain cells (top) and redistribution cells (bottom). Shown are population distributions of Pearson correlation of HD-tuning scaling shape that were computed for two separate parts of the behavioral sessions (odd versus even quartiles). **d**, Same for stability of AV-tuning. Average Pearson correlation coefficient *r* is indicated above each population histogram. Note that the shapes of HD-tuning scaling were stable for both gain and redistribution cells (c – distributions are skewed towards high correlation values), whereas the shapes of AV-tuning were stable only for gain cells (d-top) but not for redistribution cells (d-bottom). This finding was consistent with the observations that, first, for redistribution cells the firing-rate does not change with angular velocity, and therefore any fluctuations in the firing-rate of these neurons as a function of angular-velocity (i.e. AV-tuning) are likely random (hence the absence of stability as reflected by nonsignificant correlations in panel d-bottom); and second, for gain cells, HD-tuning scaling by angular velocity occurred via addition of spikes at the preferred HD, resulting in an accompanying AV-tuning of these neurons (hence the high correlation values in panel d-top).

**Supplementary Fig. 5.**
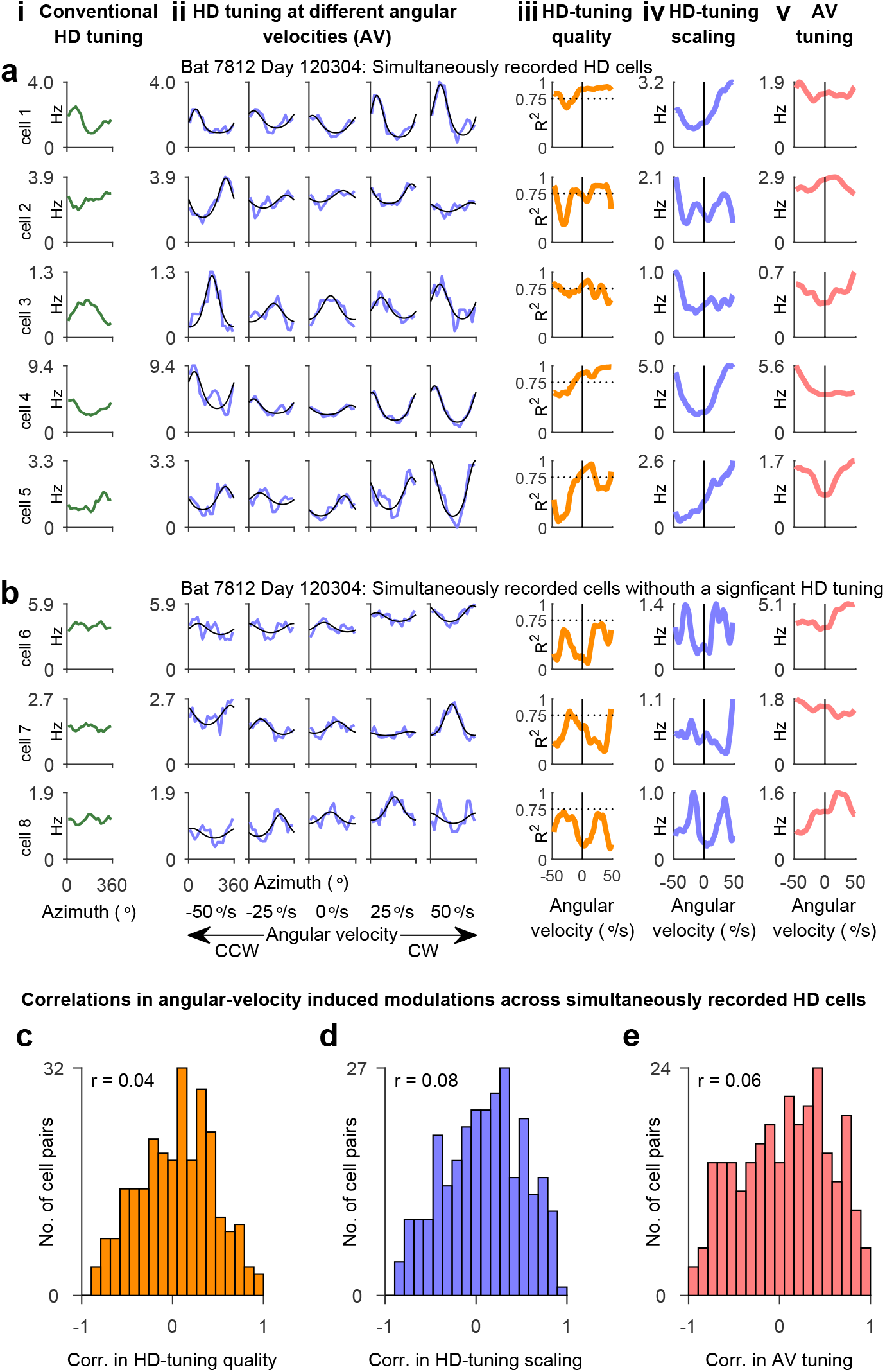
Heterogeneous modulations by angular velocity across simultaneously recorded cells. **a-b**, Examples of a single session with 8 simultaneously recorded cells – 5 HD cells (a) and 3 non HD cells (b). Columns: *i*, Conventional HD tuning computed for the entire session. *ii*, HD tuning curves (blue) computed for epochs with different angular velocities, overlaid with von Mises fits (black). *iii*, Quality of HD tuning as function of angular velocity. *iv*, HD-tuning scaling by angular velocity. *v*, AV tuning. **c-e**, Distributions of Pearson correlation coefficients between pairs of simultaneously-recorded cells, computed for shapes of HD-tuning quality (c), HD-tuning scaling (d), and AV-tuning (e). Histograms depict all pairs of simultaneously recorded HD cells.

**Supplementary Fig. 6.**
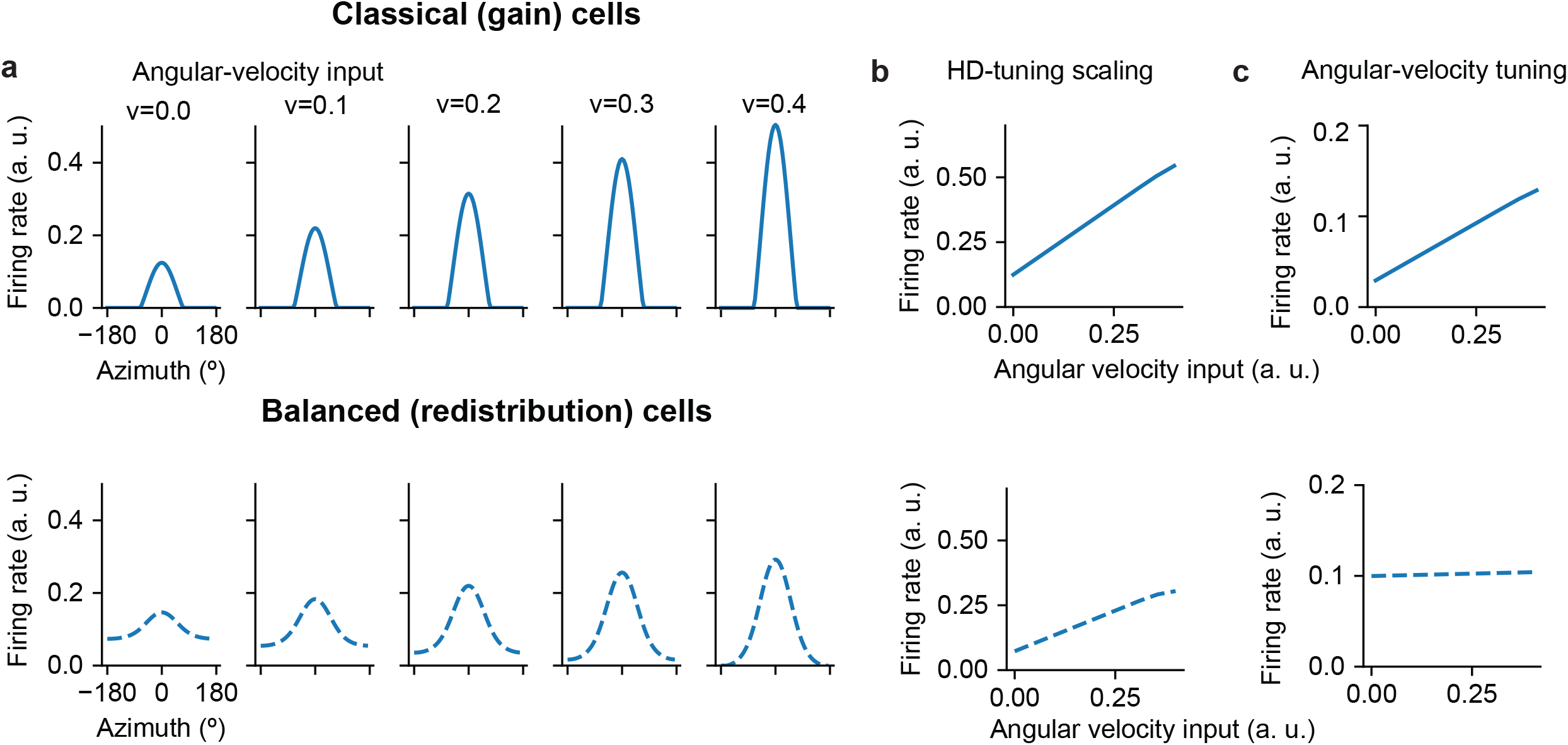
Static effective model with one ring for each classical and balanced population. Combining classical cells with balanced cells does not disrupt the existence of a localized bump of activity. Classical and balanced cells are coupled as shown in main Fig. 5b. The relationship between mean activity and angular-velocity input for the balanced cells becomes weaker with increased inhibitory feedback (Methods). **a**, Population activity profiles showing HD-tuning for different angular velocity inputs (v, columns), for classical (top) and balanced cells (bottom). **b-c**, HD-tuning scaling (b) and angular-velocity tuning (c) for classical cells (top) and balanced cells (bottom). Note that HD-tuning scaling increases with angular-velocity for both classical and balanced cells, but AV-tuning was apparent only in classical cells.

**Supplementary Fig. 7.**
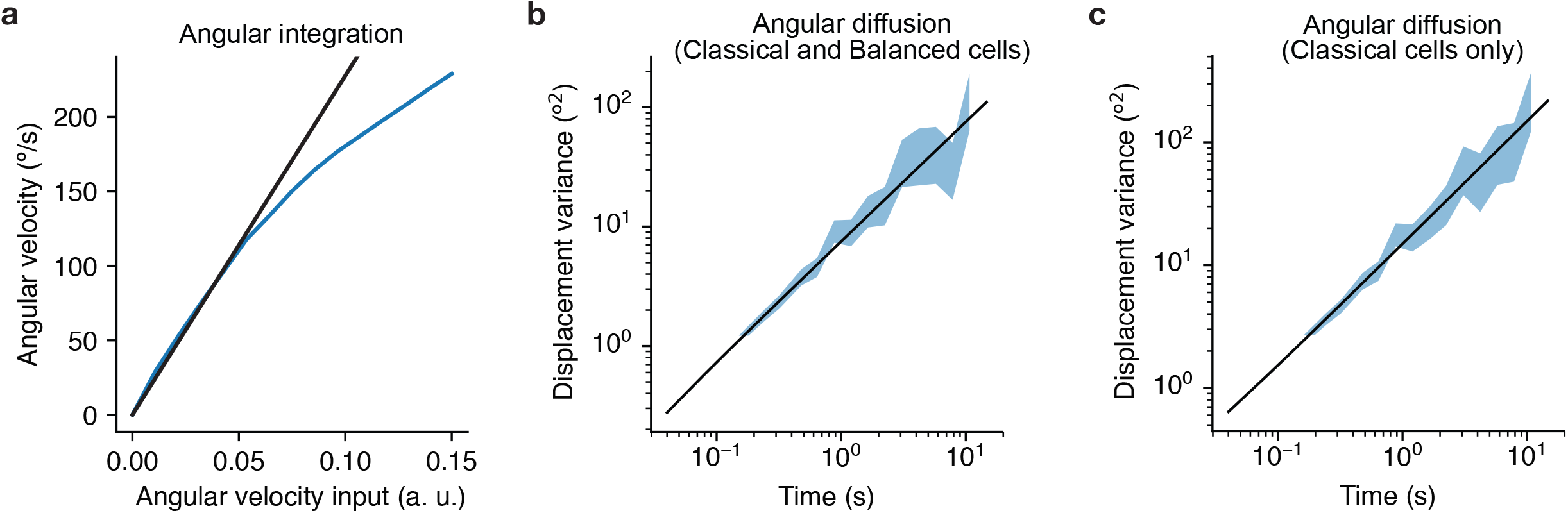
Angular path integration and bump diffusion in the presence of noisy neurons in the 6-ring model. **a**, Angular velocity of the bump of activity (blue) as a function of input angular velocity. The bump’s angular velocity changes linearly as a function of input angular velocity, covering the velocity range observed experimentally (the relationship remains linear until ~150 °/s; Black, linear fit). This indicates that combining classical (gain) cells with balanced (redistribution) cells in the 6-ring model does not disrupt angular-velocity integration. **b**, Bump diffusion in the presence of noise in the 6-ring model. Plotted is the variance of bump position as a function of time (blue; double logarithmic scale) and a linear fit (black). A small diffusion coefficient (D = 1.5 10^−3^ o2/s), defined as the intercept of the linear fit (black line), is indicating that in the 6-ring model the bump location is robust to noise. **c**, Same as b, but for classical (gain) cells only (methods). The diffusion coefficient is weakly affected (D = 3.0 10^−3^ o2/s).

**Supplementary Fig. 8.**
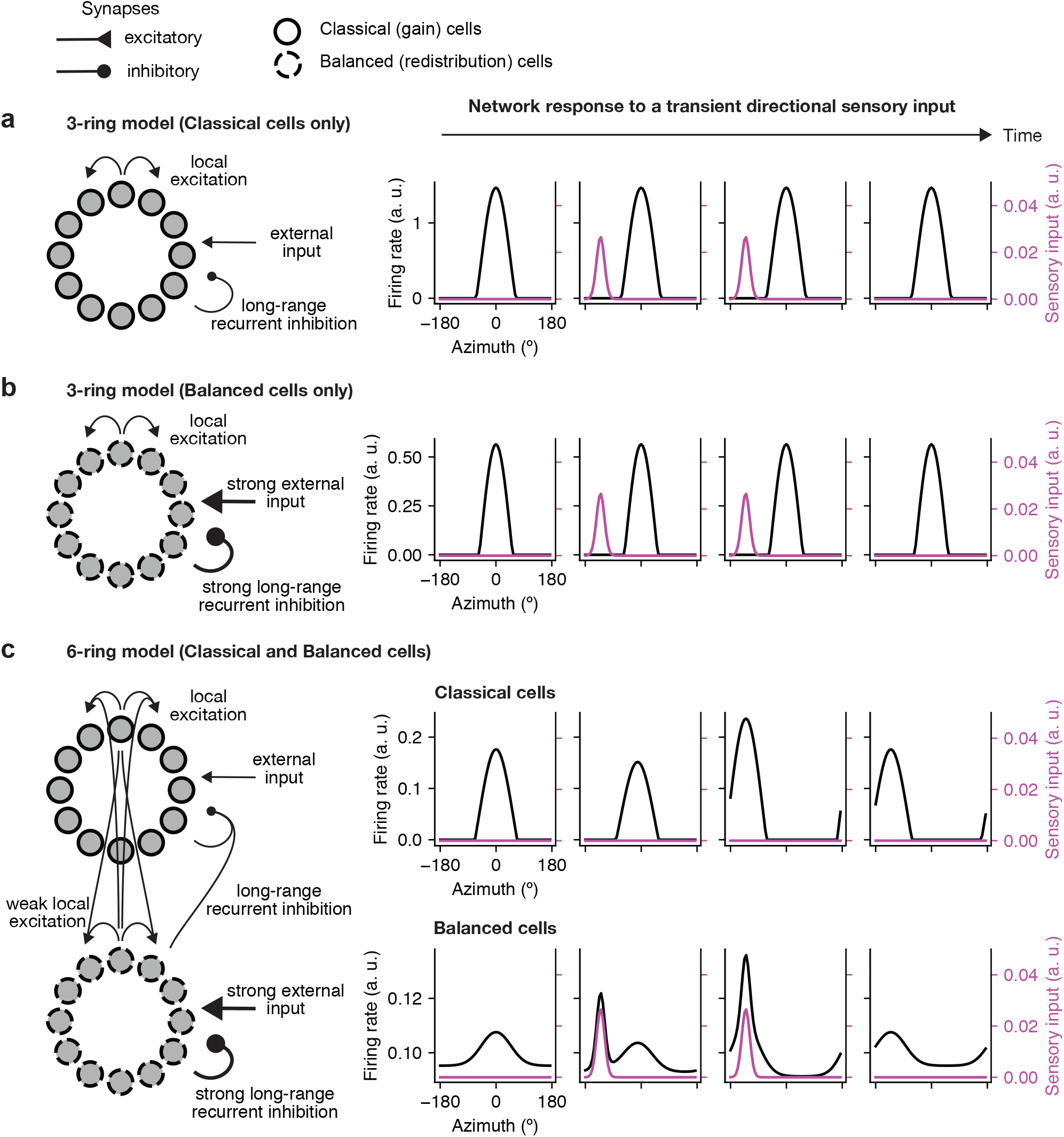
Updating the bump position by external sensory input. Network architecture schematics (left) and bump dynamics (right panels, black curves), in response to the presentation of a transient directional sensory input. A weak and narrow input was transiently delivered to the network (magenta). Time increases from left to right panels. **a**, 3-ring model composed of classical cells only. The sensory input did not cause a movement of the bump, due to strong inhibition outside the preferred direction of classical cells. Therefore, sensory inputs that are not sufficiently strong to overcome this inhibition would not be able to update the bump position. **b**, 3-ring model composed of balanced cells only. Sensory inputs did not produce a bump movement in this network as well, because generation of a localized and persistent bump requires the presence of strong suppression of firing in the non-preferred direction. Hence, a network of balanced cells exhibited a behavior similar to a network composed of classical cells only (a). **c**, 6-ring model composed of both classical (top row) and balanced cells (bottom row). The addition of balanced cells operating in the supra-threshold regime to classical cells, increased the sensitivity of the network to weak and localized sensory inputs. In this 6-ring network, the appearance of a sensory input led to a shift of the bump position towards the direction of the sensory input. The update of the bump position occurred first in balanced cells and then propagated to the classical cells via recurrent interaction.

**Supplementary Fig. 9.**
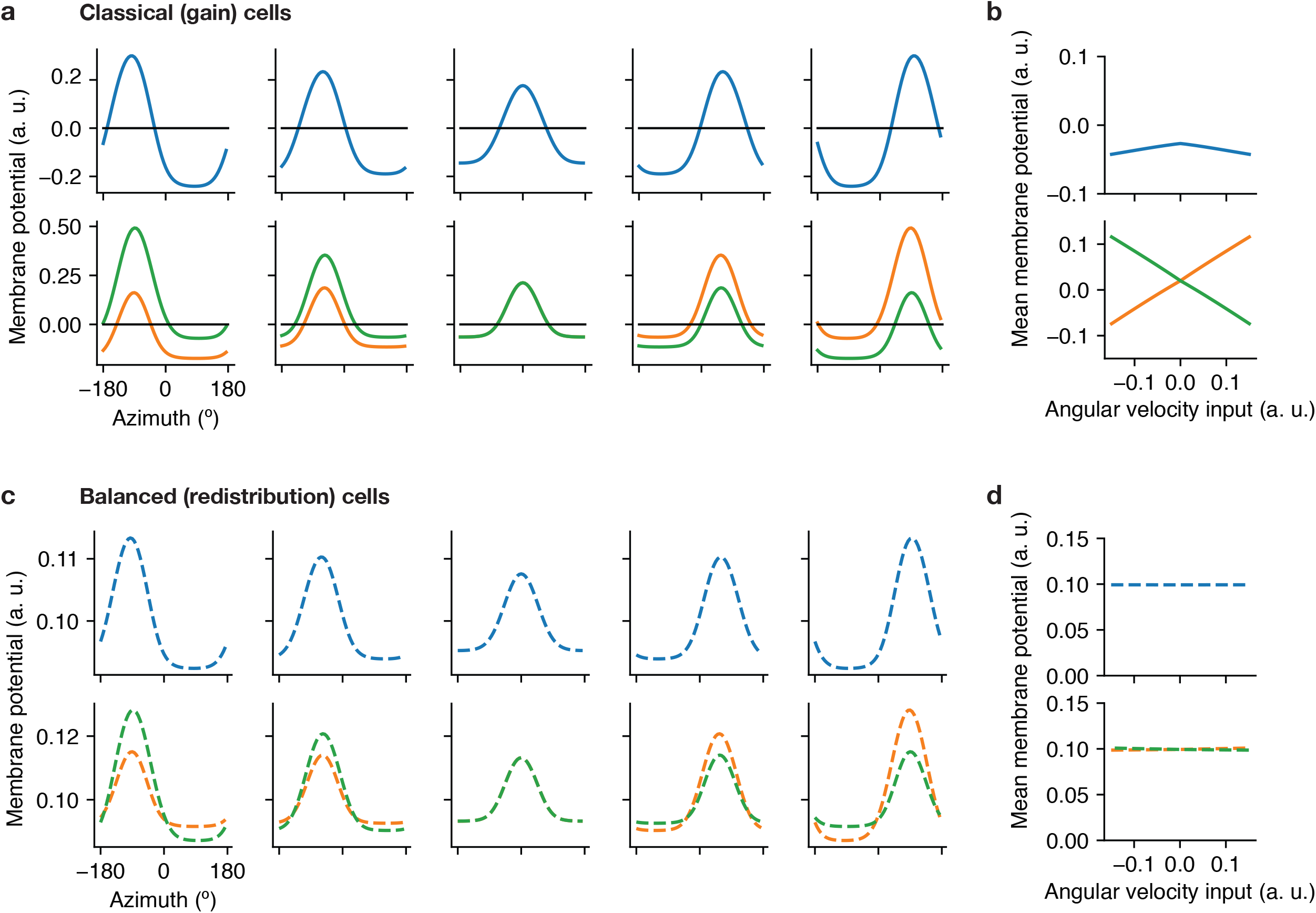
Model predictions for the membrane potential dynamics of gain and redistribution cells. **a**, HD tuning of the membrane potential at varying angular velocity for classical (gain) neurons on different rings (blue – central ring, green – left ring, orange-right ring). **b**, Mean membrane potential as a function of angular velocity. **c-d**, Same as (a-b), for balanced (redistribution) neurons. Note the difference in the membrane potential profiles for gain and redistribution cells (compare b and d).

